# MSGFLEX: An optimized end-to-end pipeline for proteomics and metaproteomics analysis built on MS-GF+ with a dedicated rescoring model, XPECTRA

**DOI:** 10.64898/2026.11.19.767922

**Authors:** Tulasi Rao Relangi, Harrison A. Hall, Aaron T. Wright

## Abstract

Mass spectrometry-based proteomics workflows often face competing demands between identification sensitivity, false discovery rate (FDR) control, and computational scalability, particularly for metaproteomic datasets that require searching large and complex sequence databases. Here, we present MSGFLEX, an integrated end-to-end workflow that combines database searching, machine-learning-based rescoring, and peptide-to-protein quantitative inference. MSGFLEX integrates three components: SPARX handles raw file conversion, and MS-GF+ database searching; XPECTRA is a cross-validated XGBoost ensemble that rescores peptide-spectrum matches by combining ensemble predictions with MS-GF+ search evidence through noisy-OR fusion model. QUANTIX provides protein quantification through multiple protein rollup strategies. Across four benchmarking datasets, XPECTRA consistently increased peptide and protein identifications relative to Percolator, MS^2^Rescore, and MSBooster. On a human gut metaproteome dataset, XPECTRA achieved the largest effect size among all methods evaluated (Cohen’s *d* = 0.95; *p* < 0.0005). SHapley Additive exPlanations (SHAP)-based analysis indicated that XPECTRA learned consistent relationships from spectral and physicochemical features. Database partitioning does not introduce detectable bias in peptide identification or FDR estimation, while QUANTIX’s *DWrollup* provides the most accurate recovery of known protein abundance changes in a ground-truth spike-in benchmark. Together, these results establish MSGFLEX as a scalable and interpretable framework for mass-spectrometry based proteomics and metaproteomics.

## 1 Introduction

Shotgun proteomics data analysis is a multi-step process in which raw mass spectra are searched against a protein sequence database, peptide-spectrum matches (PSMs) are filtered according to confidence criteria, and surviving peptide identifications are quantified across samples. Although these stages are typically performed sequentially, they are often handled by separate software tools rather than a unified analytical framework[1]. As a result, proteomics workflows frequently require substantial manual coordination across multiple processing steps before biologically interpretable results can be obtained.

Database search engines generate candidate peptide identifications, whereas machine learning-based rescoring approaches such as Percolator [2, 3] and related methods improve PSM discrimination by learning data-driven scoring functions that complement and reweight the evidence provided by database search engines[4]. Similarly, specialized tools have been developed for downstream quantitative analysis, enabling peptide and protein abundance estimation across experimental samples. Collectively, these advances have substantially improved proteomic identification and quantification capabilities.

Despite these advances, most tools remain developed, validated, and deployed as standalone components. Consequently, evaluations of modern rescoring methods often focus primarily on identification yield, whereas their downstream effects on quantitative proteomic analyses are less commonly examined. The need to connect multiple independent tools introduces practical challenges related to file-format compatibility, differences in how confidence metrics and false discovery rates (FDRs) are estimated and reported, and overall workflow reproducibility. These limitations become particularly pronounced in metaproteomics, where complex microbial communities are commonly analyzed against large, highly redundant multi-organism sequence databases. Such database search spaces can reduce identification sensitivity and have motivated the development of strategies such as database partitioning, advanced PSM rescoring, and quantification approaches designed to tolerate sparse identifications [5, 6]. Beyond their effects on peptide identification, these challenges can propagate to downstream stages of analysis, influencing PSM discrimination, protein inference, and quantitative protein abundance estimates. This dependency highlights the need for integrated workflows that enable the consequences of upstream analytical decisions to be assessed throughout the entire metaproteomics pipeline.

To address these limitations, MSGFLEX was developed as an integrated end-to-end workflow that combines database searching, machine learning-based PSM rescoring, and quantitative proteomics within a single analytical framework. The goal is to reduce workflow fragmentation while providing a reproducible path from raw mass spectra to quantitative protein abundance profiles.

MSGFLEX consists of three tightly integrated modules. First, SPARX performs raw-file conversion and MS-GF+-based database searching, with optional database splitting and binning strategies for large and complex sequence databases. Second, XPECTRA applies a leakage-controlled, cross-validated XGBoost-based ensemble model that integrates complementary spectral, sequence, and physicochemical features to improve PSM confidence. Third, QUANTIX provides flexible peptide-to-protein relative quantification through multiple peptide-to-protein rollup strategies. Unlike conventional evaluations of PSM rescoring tools, XPECTRA is assessed not only by identification yield at controlled FDR thresholds but also by its impact on downstream quantitative proteomic measurements. Its performance is benchmarked against three established rescoring approaches, Percolator, MS^2^Rescore, and MSBooster [2, 7, 8], using public datasets spanning synthetic communities, environmental and host-associated microbiomes, and human clinical samples. Together, these analyses demonstrate that MSGFLEX provides a unified and reproducible workflow from raw mass spectra to confidently filtered quantitative protein profiles.

## 2 Results

### 2.1 MSGFLEX architecture

MSGFLEX comprises three sequential modules that provide an integrated end-to-end workflow from raw mass spectra to quantitative protein profiles. Intermediate results are exchanged through standardized tabular formats, enabling seamless propagation of identification and confidence information between analysis stages. This design eliminates manual reconciliation across disparate tools and provides a unified framework for evaluating the effects of database searching, rescoring, and confidence filtering on downstream quantitative results (Fig. 1).

**Fig. 1.**
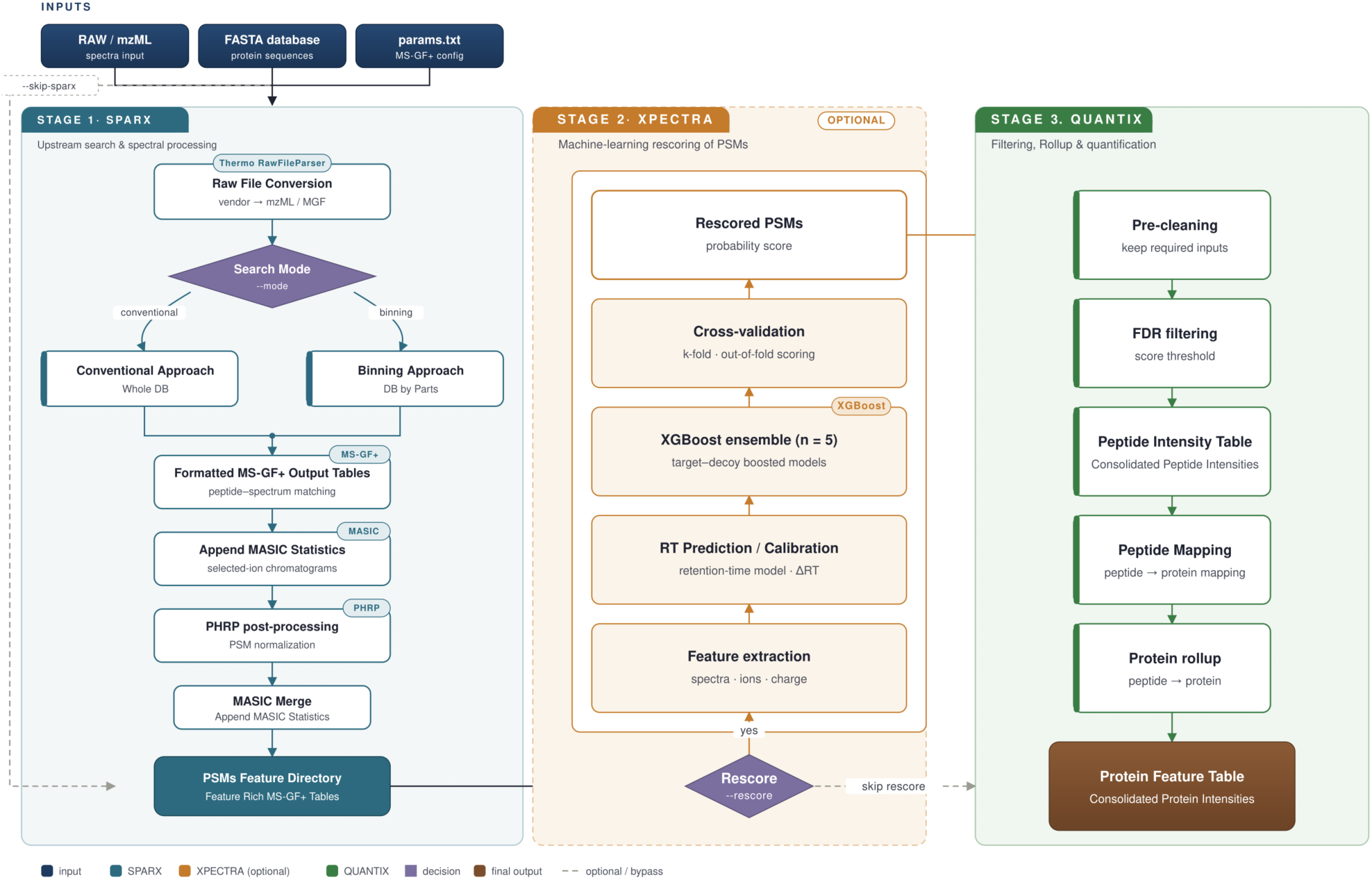
Overview of the MSGFLEX pipeline architecture. MSGFLEX processes raw mass spectrometry data through three sequential stages. **Stage 1 (SPARX)** file conversion, MS-GF+ database search in whole-database or binning (database split into parts) mode, and PSM post-processing. **Stage 2 (XPECTRA, optional)** an optimized machine-learning rescoring of PSMs. **Stage 3 (QUANTIX)** filters PSMs by score threshold to control FDR, consolidates peptide-level intensities, and performs protein rollup to produce a final consolidated protein feature table. Pipeline behavior is configured via params.txt; the –skip-sparx flag allows entry directly at the PSM feature stage for pre-processed inputs. Colors indicate module membership (input, teal: SPARX, orange: XPECTRA, green: QUANTIX, purple: decision points, brown: final output); dashed arrows indicate optional or bypass paths.

Each module can be used independently or as part of the complete MSGFLEX workflow. SPARX output can be passed directly to QUANTIX for quantification without rescoring. Alternatively, previously generated SPARX PSM tables can be supplied directly to XPECTRA using the –skip-sparx option, allowing rescoring to be performed without repeating the database search step. Consequently, MSGFLEX can be deployed either as a complete raw-to-quantification end-to-end pipeline or as modular rescoring or quantification component within existing proteomics workflows. The pipeline is supported on Linux and can be executed through either a command-line or an optional graphical user interface (Usage of MSGFLEX Command Line Interface/Graphical User Interface depicted in Supplementary Methods 1; Code Availability).

A key design goal of SPARX was to reduce the substantial wall-clock time that large-cohort MS-GF+ searches impose in practice. To this end, SPARX incorporates NUMA-aware CPU allocation and dynamic thread management that right-sizes the number of MS-GF+ tasks per sample at runtime. To quantify this benefit, 16 randomly selected samples from the PRIDE dataset PXD017035[9] were searched against the SIHUMIx reference database. On a workstation with 128 GB RAM and 48 threads, task optimization reduced the per-sample task count from 144 to 8 and cut search times 2.3–2.4-fold on average relative to the default MS-GF+ configuration (paired Wilcoxon signed-rank test, *p <* 0.0001; Fig. S1a). Within the database-binning workflow, the same scheduling produced a similar benefit: three-bin partitioning reduced search time 1.7–1.8-fold relative to the default configuration (paired Wilcoxon signed-rank test, *p <* 0.0001; Fig. S1b).

### 2.2 XPECTRA rescoring improves PSM and peptide identification

XPECTRA was evaluated against established rescoring approaches, including Percolator[2, 3], MS^2^Rescore[7], and MSBooster[8] each used in combination with Percolator. Unlike modern rescoring approaches that frequently rely on external deep-learning models for peptide property prediction, such as Retention-Time (RT) and fragment-ion intensity, XPECTRA uses a classical machine-learning framework that integrates search-engine scores, spectral features, RT features, and precursor-level characteristics. This design reduces software dependencies and computational overhead while maintaining strong identification performance. To ensure a fair comparison, all datasets were additionally rescored with deep learning-based RT features. Protein-level gains were evaluated using the unique consensus-mapped protein groups described in the Methods section under Benchmarking against Percolator, MS^2^Rescore, and MSBooster+Percolator.

Four different benchmark datasets were used to evaluate the performance of XPECTRA and other rescoring methods. Across all four benchmark datasets, XPECTRA consistently improved identification yield over native MS-GF+ baseline at 1% FDR, with the largest gains observed at PSM and protein levels (Fig. 2, a,f). On the synthetic community dataset (PXD006118 [10]) and human gut metaproteome dataset (PXD035981[11]), XPECTRA recovered the highest number of additional PSMs of any compared model, exceeding Percoltor, MS^2^Rescore, and MSBooster+Percolator by a wide margin (+14.1% and +26.6% netgain, respectively, versus the next-best compared model MSBooster+Percolator at +8.4% and +25.7%, Fig. 2, d). On the marine metaproteome (PXD052873[12]), a similarly large lead was observed (+48.3% versus +41.6% for MSBooster, Fig. 2, d).

**Fig. 2.**
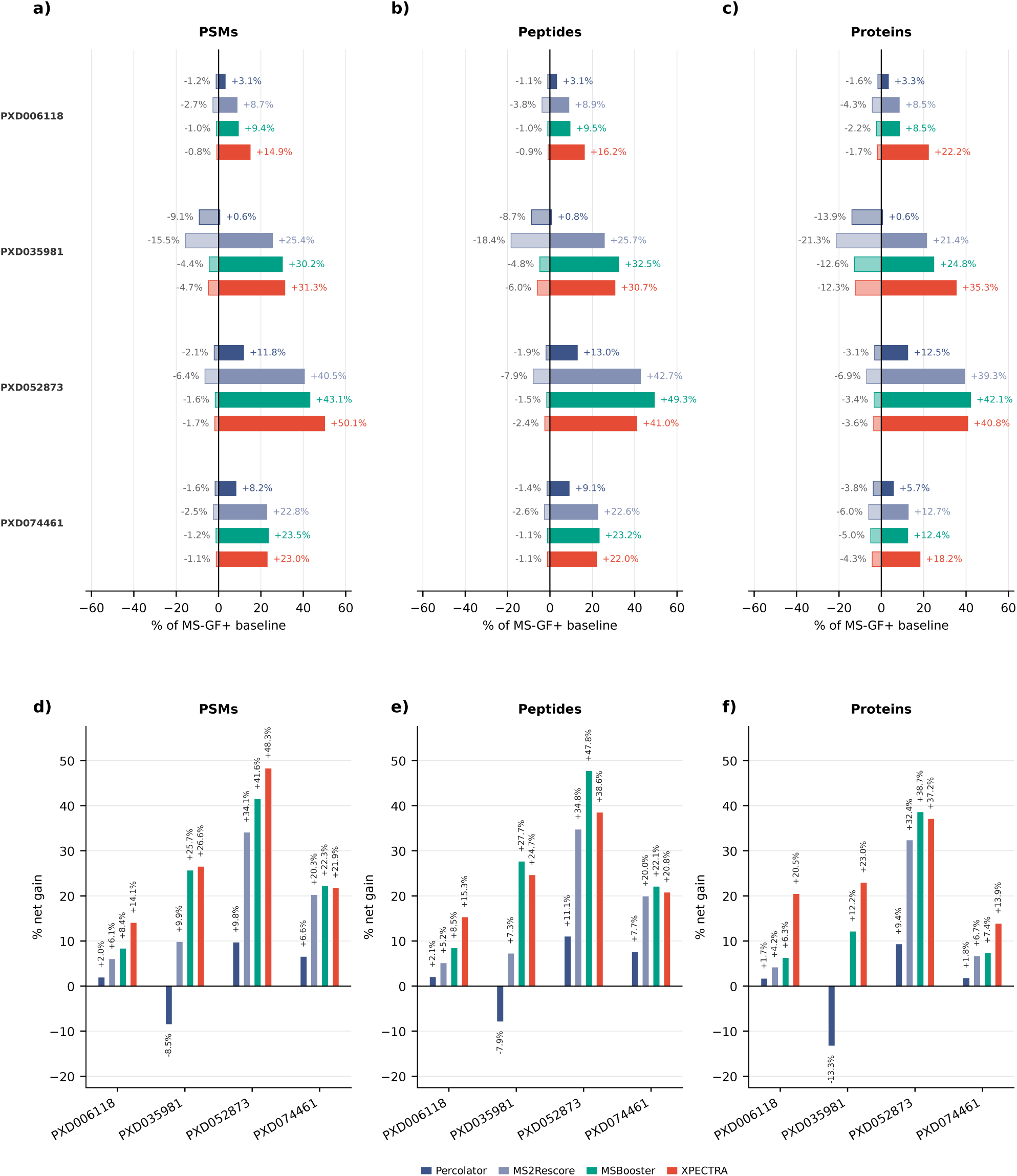
XPECTRA rescoring improves identification yield across four benchmark datasets relative to MS-GF+ baseline and comparator tools. **a–c** Percentage change in identified PSMs (**a**), peptides (**b**), and proteins (**c**) relative to the MS-GF+ baseline (0%, vertical line) for each of four comparator tools (Percolator, MS^2^Rescore, MSBooster, XPECTRA) across four benchmark datasets (PXD006118, PXD035981, PXD052873, PXD074461). For each tool/dataset pair, losses (targets removed relative to baseline) are shown to the left of the axis and net gains (targets added) to the right; XPECTRA (red) consistently achieves the smallest losses and largest gains across all three identification levels. **d–f** Net percentage gain (gains minus losses) in PSMs (**d**), peptides (**e**), and proteins (**f**) for each tool across the same four datasets, summarizing the trade-off shown in **a–c** into a single net metric per tool/dataset combination. XPECTRA achieves the highest net gain in the majority of tool/dataset comparisons, with the exception of PXD052873, where MSBooster shows a marginally higher net gain at the protein level. Percentages are computed relative to MSGF+ baseline identification counts at 1% FDR for each dataset.

At the peptide level, XPECTRA and MSBooster produced comparable gains across the human gut, marine, and human datasets, with net peptide-level improvements differing by only a few percentage points (Fig. 2b,e). This pattern likely reflects the redundancy of peptide evidence in complex datasets, where multiple PSMs frequently map to the same peptide sequence, thereby attenuating the translation of PSM-level gains into increases in unique peptide identifications. Notably, on the synthetic community dataset, where the underlying peptide composition is well defined, XPECTRA again showed a clear advantage (+15.3% versus +8.5% for MSBooster), consistent with its strong PSM-level performance.

Protein-level recovery showed the clearest overall advantage for XPECTRA, which achieved the highest net protein gain in three of the four datasets (synthetic: +20.5%; human gut: +23.0%; human: +13.9%), each exceeding the next-best comparator by a substantial margin. On the marine metaproteome, XPECTRA’s protein-level gain (+37.2%) was essentially matched by MSBooster (+38.7%), reflecting the same convergence observed at the peptide level for this dataset (Fig. 2, c&d). Taken together, these results indicate that while XPECTRA’s peptide-level gains are broadly comparable to MSBooster’s on complex, real-world samples, its improvements consistently translate into superior or matched protein-level recovery across all four benchmark datasets – the identification tier most directly relevant to downstream biological interpretation.

To assess whether XPECTRA’s identification gains reflected genuinely higher-confidence PSMs rather than a relaxed decision boundary, we compared the separation between target and decoy score distributions across all methods using Cohen’s *d* effect size on the human gut metaproteome (PXD035981; Fig. 3). The unrescored MS-GF+ baseline and MSGFLEX-SPARX showed near identical target-decoy separation (*d* = 0.70 for both), consistent with SPARX’s PHRP standardization preserving the native MS-GF+ scoring distribution (Fig. 3a). Percolator produced a modest improvement (*d* = 0.73, Fig. 3b), while MS^2^Rescore and MSBooster showed progressively greater separation (*d* = 0.79 and *d* = 0.91, respectively), each introducing a visibly bimodal target distribution – a low-scoring mode overlapping the decoy distribution alongside a well seperated high-confidence mode (Fig. 3d,e). XPECTRA achieved the largest seperation of any method (*d* = 0.95, Fig. 3f). To confirm this ranking was not an artifact of sampling variability, we performed a bootstrap resampling analysis (2,000 resamples per method) comparing XPECTRA’s effect size against each of the other models. XPECTRA’s separation was significantly greater than all five other methods (all bootstrap 95% CIs on the difference excluding zero, *p <* 0.0005 in each case) including closest competing method MSBooster+Percolator (Δ*d* = 0.034, 95% CI[0.032, 0.035]). Effect sizes remained unchanged between the baseline and SPARX workflows, increased modestly for the comparator methods, and increased most strongly for XPECTRA. These results indicate that XPECTRA enhances target-decoy discrimination while simultaneously increasing identification yield, suggesting improved scoring performance across datasets.

**Fig. 3.**
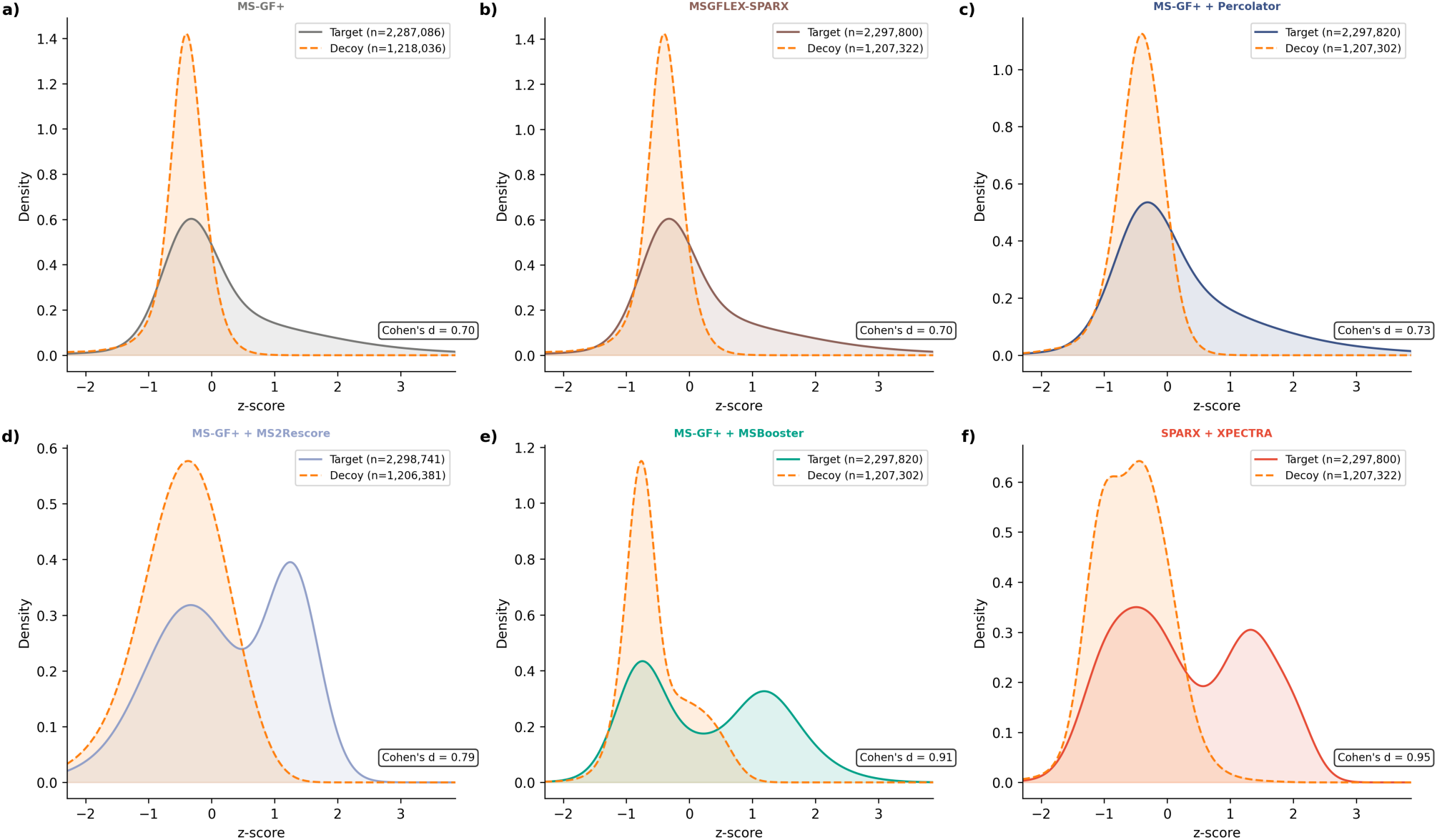
Target-decoy score separation across search engines and rescoring tools. Kernel density estimates of target (solid) and decoy (dashed) score distributions, converted to z-scores for cross-panel comparability, for (**a**) MS-GF+ baseline, (**b**) MSGFLEX-SPARX, (**c**) MS-GF+ with Percolator rescoring, (**d**) MS-GF+ with MS^2^Rescore, (**e**) MS-GF+ with MSBooster, and **f** SPARX with XPECTRA rescoring, evaluated on PXD035981. Target and decoy counts (*n*) are shown per panel. Effect size (Cohen’s *d*) quantifies the separation between target and decoy score distributions; XPECTRA achieves the largest separation (Cohen’s *d* = 0.95), exceeding all comparators (bootstrap (*n* = 2000) resamples, *p <* 0.0005 for all pairwise comparisons). The bimodal target distributions visible in **d–f** reflect a subpopulation of high-confidence rescued targets that are well-separated from the decoy distribution, a pattern absent in the unimodal baseline and Percolator distributions (**a–c**).

To investigate the basis of XPECTRA’s identification gains, RT behavior was examined for PSMs gained by XPECTRA in a representative sample (Fig. S2a–d). Following RT prediction and calibration, gained PSMs closely followed the predicted-versus-observed RT relationship and exhibited residual distributions similar to those of commonly identified PSMs, consistent with expected chromatographic behavior and indicating that the gains are not artifacts of rescoring. Comparison of native MS-GF+ SpecEValue scores with XPECTRA rescoring scores in the human gut metaproteome dataset PXD035981 revealed a substantial *rescue region* comprising PSMs that failed the MS-GF+ 1% FDR threshold but passed the XPECTRA 1% FDR threshold (Fig. S2e). This region was dominated by target PSMs (231,696 targets versus 7,052 decoys), indicating that the recovered identifications are enriched for correct matches, while the global 1% FDR remained controlled.

XPECTRA improves identification sensitivity by reclassifying borderline PSMs that carry weak search-engine evidence but retain sufficient discriminative signal across the broader spectral and physicochemical feature space used for rescoring. To assess whether this recovery was biased toward particular functional categories, COG (Clusters of Orthologous Groups) classification was performed on proteins uniquely gained, shared, and lost relative to the MS-GF+ baseline. No significant enrichment or depletion was detected for any COG category (Fig. S3), and the recovered proteins were distributed broadly across functional classes — confirming that the identification gains reflect a general increase in sensitivity rather than preferential recovery of specific protein groups.

### 2.3 Feature attribution reveals rescoring drivers

To characterize the features driving XPECTRA predictions, we computed SHAP (SHapley Additive exPlanations) values for each PSM using TreeExplainer[13, 14] and averaged them across the five ensemble models used within each cross-validation fold on human gut metaproteome dataset (PXD035981). Because the final XPECTRA score is generated by combining the ensemble prediction with native MS-GF+ evidence through a post hoc noisy-OR fusion step[15], SHAP analyses were performed on the pre-fusion ensemble prediction (rescore_prob), where feature attributions remain additive and directly interpretable.

The peptide-level best spectral E-value, pep_best_specev, was the most important feature according to both mean absolute SHAP value and XGBoost gain, consistent with its role as the strongest source of search-engine evidence (Fig. 4a). Matched-ion count features (num_matched_ions_total and num_matched_y_ions) and the transformed spectral E-value (neg_log_specEvalue) also ranked highly by both metrics, indicating that XPECTRA relies heavily on direct spectral-match evidence (Fig. 4a). RT match probability (rt_match_prob) ranked more highly by XGBoost gain than by mean SHAP value, suggesting frequent use in tree splitting but a more modest contribution to final prediction confidence.

**Fig. 4.**
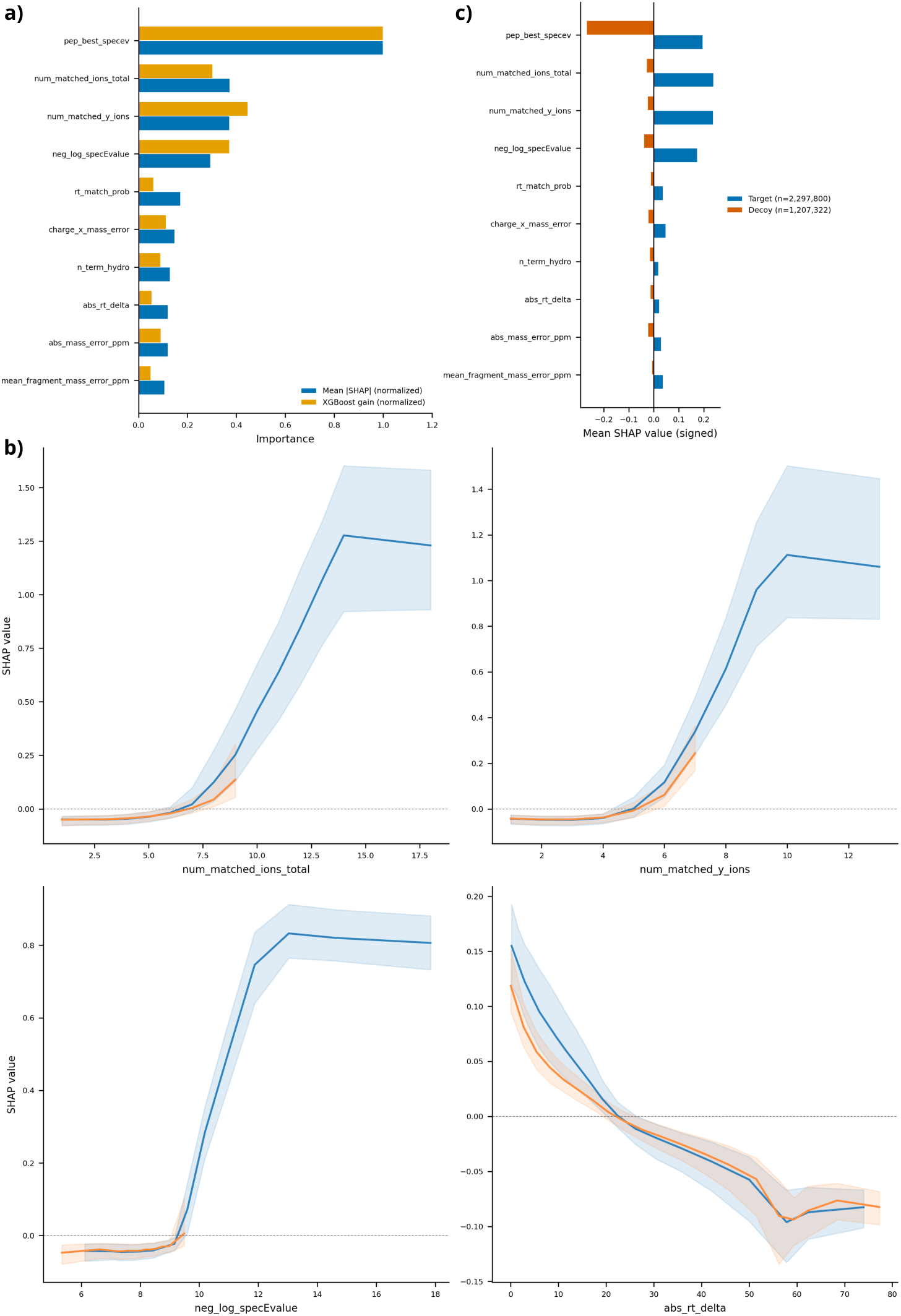
XPECTRA learns discriminative spectral and physicochemical features. (**a**) Feature importance ranking comparing mean absolute SHAP value (blue) against XGBoost gain (gold), both normalized to their respective maxima, across the ten features used by the XPECTRA rescoring model. Four features (pep_best_specev, num_matched_ions_total, num_matched_y_ions, neg_log_specEvalue) rank highly by both metrics; SHAP additionally identifies abs_rt_delta and fragment mass error metrics as informative despite comparatively low XGBoost gain, consistent with known limitations of gain-based importance under feature correlation. (**b**) SHAP dependence plots for the four highest-impact continuous features (num_matched_ions_total, num_matched_y_ions, neg_log_specEvalue, abs_rt_delta), showing mean SHAP value (line) and 95% confidence interval (shaded band) as a function of feature value, stratified by target (blue) and decoy (orange) PSM assignment. (**c**) Mean signed SHAP value per feature, split by target (blue, (*n* = 2, 197, 800)) and decoy (orange, (*n* = 1, 207, 322)) PSMs, showing the direction and magnitude of each feature’s contribution to the rescoring probability. All panels were computed on PXD035981 using TreeExplainer.

To assess whether feature contributions differed between target and decoy PSMs, SHAP dependence profiles of the most influential features were examined separately for each class (Fig. 4b). Matched-ion count and spectral E-value features diverged sharply between classes: SHAP values for target PSMs rose steeply with increasing num_matched_ions_total, num_matched_y_ions, and neg_log_specEvalue, whereas decoy PSMs showed a comparatively muted response across the same feature range, with the divergence particularly pronounced for neg_log_specEvalue (Fig. 4b). This pattern indicates that XPECTRA learned target-specific relationships between spectral evidence and identification confidence. In contrast, abs_rt_delta showed near-identical trends for both target and decoy PSMs, consistent with its role as a general quality indicator rather than a primary target-decoy discriminator.

Consistent with the dependence-profile analysis, directonal mean SHAP values were systematically positive for target PSMs and negative for decoy PSMs across all topranked features (Fig. 4c), indicating the model consistently pushed predictions toward higher confidence for targets and lower confidence for decoys regardless of feature identity. This effect was most pronounced for pep_best_specev, which showed the largest magnitude divergence between target and decoy classes (decoy mean SHAP ≈ −0.27 vs. target mean SHAP ≈ +0.20 among *n* = 1, 207, 322 decoy and *n* = 2, 297, 800 target PSMs), consistent with its role as the primary discriminative feature identified in the global importance ranking. Matched-ion count and spectral E-value features showed the same directional pattern at proportionally smaller magnitude. The consistency of this sign inversion across features with otherwise widely varying overall importance indicates the model’s target/decoy discrimination is not driven by a small number of dominant features acting in isolation, but reflects a coherent directional signal distributed across the feature set.

Pooled (non-class-stratified) feature-importance rankings and SHAP dependence profiles are provided in Fig. S4. Raw signed mean SHAP magnitudes reproduced the same overall importance hierarchy observed in Fig. 4a, with num_matched_ions_total, num_matched_y_ions, and neg_log_specEvalue showing the largest average contributions to model output. Likewise, pooled dependence profiles showed monotonic trends consistent with the class-stratified relationships in Fig. 4b, indicating that the target-specific divergence arises from stratification of a broader population-level pattern rather than from the stratification procedure itself.

### 2.4 Binning/splitting large databases does not bias FDR or peptide identification

A common concern with database-splitting or database-reduction strategies is their potential impact on false discovery rate (FDR) estimation. Prior two-step approaches construct a reduced search database from preliminary identifications and subsequently re-search the same spectra against this data-derived database, introducing a dependency between database construction and peptide identification that may affect error-rate estimation and increase false-positive identifications[5, 16]. Database size can also influence target-decoy competition and, consequently, FDR estimation [17]. In contrast, our binning approach randomly splits the sequence database into non-overlapping bins before any database search is performed. Because the E-value is database-size dependent, bin-specific E-values were first corrected according to the relative sizes of the full and partitioned databases using Eq. (1) before merging the results from all bins.

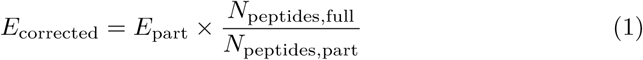

The corrected results from all bins were then merged and processed with PHRP, which recalculates spectrum- and peptide-level *q*-values from the resulting target-decoy distributions.

To evaluate whether database binning introduces bias, we compared the SPARX binned workflow (using 10 bins) against a conventional unified, non-partitioned search of the same database (223,441 sequences). Kernel density estimation (KDE) and empirical cumulative distribution functions (ECDFs) were used to visualize the distributions of search and confidence metrics, and distributional differences between split-merged and unified database searches were quantified using the Kolmogorov– Smirnov (KS) statistic. The comparison evaluated PSM- and peptide-level identification results after XPECTRA rescoring across eight randomly selected samples from the PXD057701 dataset[18]. The two workflows produced nearly identical identification results at 5% FDR, yielding 52,276 and 52,282 accepted target PSMs for the binned and unified searches, respectively. SpecEValue and EValue distributions were highly similar between the two workflows, with *KS − D* values of 0.003 and 0.112, respectively (Fig. 5a,b), indicating that database partitioning, E-value correction, and result merging did not substantially alter the overall score distributions. Consistent with this observation, the ECDFs of QValue and PepQValue showed close overlap between the two workflows, with *KS − D* values of 0.0427 & 0.0751, respectively (Fig. 5c,d), indicating that target-decoy-based confidence estimation was largely preserved following result merging.

**Fig. 5.**
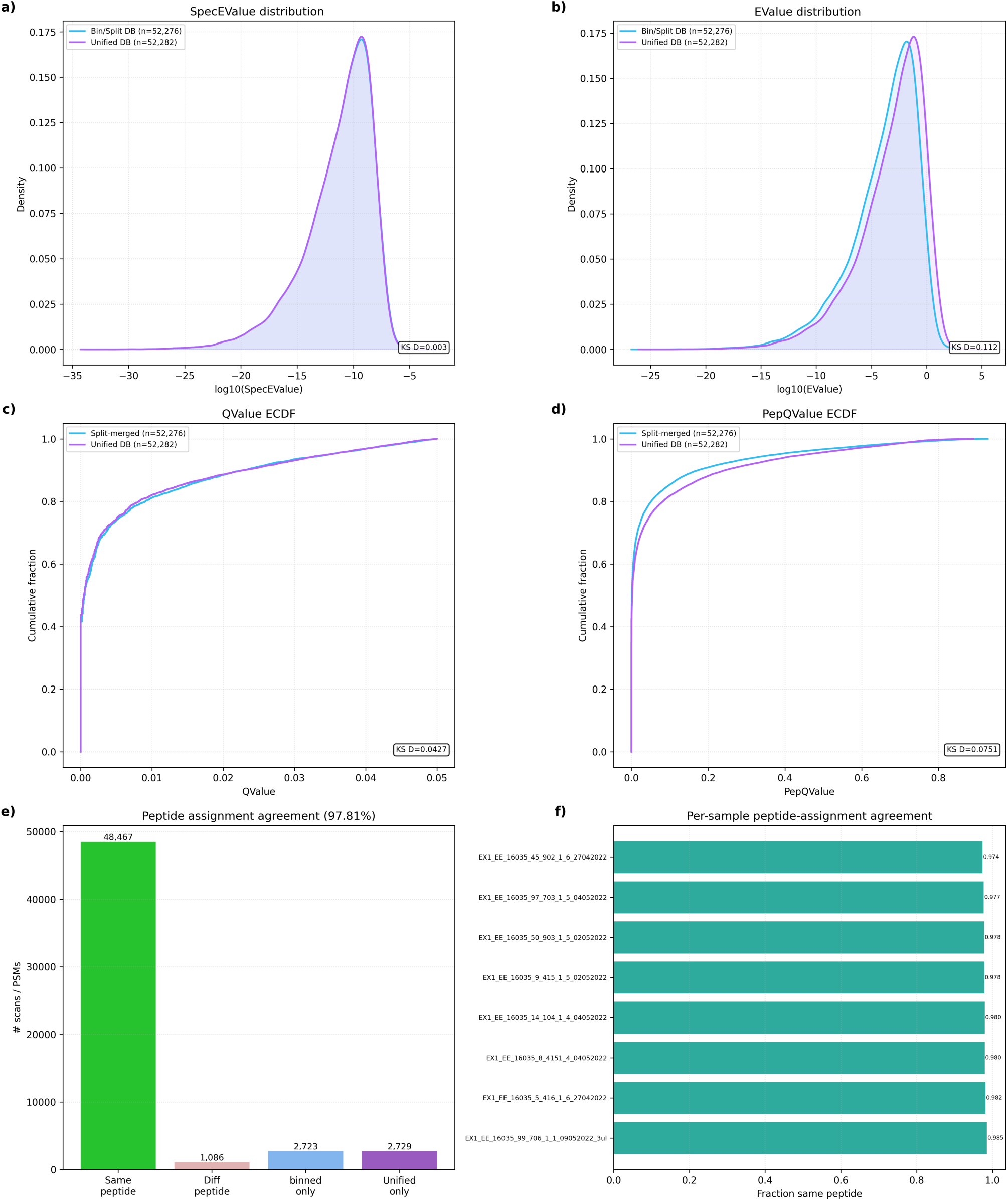
Binning does not bias identification outcomes relative to a unified database search following XPECTRA rescoring. **a, b** Kernel density estimates of log_10_(SpecEValue) (**a**) and log_10_(EValue) (**b**) distributions for PSMs identified using the binned/split database (cyan) versus the unified database (purple), showing near-identical score distributions between the two approaches. **c, d** Empirical cumulative distribution functions of QValue (**c**) and PepQValue (**d**) for the binned/split (cyan) versus unified (purple) database searches. The maximum empirical differences were small (QValue: *D* = 0.0427; PepQValue: *D* = 0.0751), indicating that the practical distributional shift was limited given the large PSM counts involved. **e** Peptide-level assignment agreement between the binned and unified approaches: the large majority of scans/PSMs (48,467) are assigned the same peptide in both approaches, with a minority assigned different peptides (1,086) or identified uniquely by one approach (2,723 split-only; 2,729 unified-only), yielding an overall peptide-assignment agreement of 97.81%. **f** Per-sample peptide-assignment agreement across eight individual samples from PXD057701, showing consistently high agreement (fraction same peptide range: 0.974–0.985) across samples. All results shown reflect PSMs after XPECTRA rescoring.

At the peptide level, 48,467 of 52,276 PSMs (97.81%) were assigned the same peptide sequence in both workflows, with per-sample agreement ranging from 97.4% to 98.5% (Fig. 5e,f). Only 1,086 PSMs were assigned different peptide identities, and the numbers of split-only and unified-only identifications were nearly identical (2,723 and 2,729, respectively). Collectively, these results demonstrate that database partitioning does not introduce detectable bias into rescored identification results and preserves score distributions, confidence estimates, and peptide assignments relative to a conventional unified database search.

Prior to XPECTRA rescoring, database partitioning increased identification sensitivity relative to a conventional unified database search. Across the same eight PXD057701 samples, the binned workflow produced 44,473 accepted target PSMs compared with 40,316 target PSMs from the unified search, corresponding to an increase of 4,157 PSMs (Fig. S5 a,b). Despite this increase in identification yield, SpecEValue, EValue, QValue, and PepQValue distributions remained highly similar between the two workflows with *KS − D* values of 0.079, 0.093, 0.0498 & 0.0475 respectively, indicating that database partitioning did not substantially alter score calibration or FDR estimation. Peptide-level agreement also remained high, with 39,498 PSMs (97.98%) assigned the same peptide sequence in both workflows. Only 814 PSMs were assigned different peptide identities. Notably, 4,161 identifications were unique to the binned workflow, whereas only four identifications were unique to the unified search (Fig. S5e). These results indicate that database partitioning improves identification recovery before rescoring while preserving score distributions and confidence estimates.

We next evaluated the impact of XPECTRA rescoring on identification yield for both the binned and unified database-search workflows (Fig. S6). XPECTRA consistently increased the number of accepted target PSMs across all samples regardless of the underlying search strategy. For the binned workflow, accepted target PSMs increased from 44,473 to 52,276, representing an increase of 7,803 PSMs. Similarly, the unified workflow increased from 40,316 to 52,282 accepted target PSMs, corresponding to an increase of 11,966 PSMs.

These gains were accompanied by a systematic shift toward lower *q*-value, as reflected by the upward displacement of the rescored QValue ECDFs relative to the original MS-GF+ results. This indicates that a larger fraction of PSMs satisfied a fixed FDR threshold following rescoring. Although database partitioning initially yielded more accepted identifications than a unified search, XPECTRA substantially improved identification sensitivity for both workflows, resulting in nearly identical final identification counts after rescoring. These findings suggest that XPECTRA effectively recovers high-confidence identifications regardless of the underlying search configuration while preserving consistent confidence estimation.

### 2.5 QUANTIX quantification is accurate and specific on ground-truth benchmarks

We validated QUANTIX using the PXD018408 dataset[19], consisting of a 2× versus 1× *E. coli* spike-in against a HEK293 human background. Analyses were restricted to the DDA acquisition mode, which provides a ground-truth differential abundance signal independent of biological variability. Validation was performed in two stages: an initial comparison of five protein-level rollup strategies, followed by a detailed assessment of the best-performing method at both the protein and peptide levels.

To establish the most suitable protein summarization strategy for subsequent analyses, we first compared five protein-level rollup methods (*Sum*, *Rrollup*, *Qrollup*, *Zrollup*, and *DWrollup*) using 2,686 candidate proteins identified from the XPECTRA-rescored crosstab at 1% FDR, requiring valid values in at least six of eight replicates per condition (75% coverage). Differential abundance was evaluated using Welch’s *t*-test with Benjamini-Hochberg FDR correction. Performance was assessed based on accuracy (Root Mean Square Error and mean bias relative to the expected log_2_ fold change of 1.0), specificity (false-positive rate among HEK293 background proteins at adjusted *q <* 0.05), sensitivity (direction-aware detection rate among *E. coli* spike-in proteins), and overall discriminatory performance (Area Under the ROC Curve, Fig. S7a).

The *DWrollup* achieved the best overall balance, with the highest AUC (0.949) and sensitivity (58.8%, 166/282 *E. coli* proteins detected), a false-positive rate of 0.67% (11/1,612 human proteins), and an Root Mean Square Error (RMSE) of 0.657 against the expected log_2_FC. The *Sum* rollup performed comparably (AUC = 0.945, sensitivity = 59.9%, RMSE = 0.654, FPR = 0.91%), with both methods showing a positive mean bias (+0.243 and +0.234, respectively), indicating a modest systematic over-estimation of the true 2-fold change. The *Rrollup* method showed lower sensitivity (41.3%) despite a comparable AUC (0.913) and the lowest false-positive rate among the top three methods (0.43%), with an RMSE of 0.418 and a negative mean bias (−0.256), indicating underestimation of fold-change magnitude (Fig. S7b,c).

*Qrollup* and *Zrollup* showed markedly reduced sensitivity (13.8% and 3.2%, respectively) despite very low false-positive rates (0.36% and 0.12%), reflecting a conservative bias that suppressed detection of true spike-in changes (Fig. S7d,e). This conservatism was also evident in filtering behavior: *Qrollup* and *Zrollup* retained fewer proteins overall after the coverage filter (*n* = 1,593, passed filtering cut-off, and 1,894, respectively, compared with 1,914 for both *Sum* and *DWrollup*). This effect was particularly evident for the *E. coli* spike-in proteins, where *Qrollup* retained only 232 proteins compared with 278 for *DWrollup*, indicating a more conservative filtering behavior that reduced recovery of true positive differential-abundance signals. Interpretation of *Zrollup* RMSE values is also less straightforward because protein abundances are represented on a standardized *z*-score scale rather than the original log_2_ abundance scale. Based on this comparison, the *DWrollup* which achieved the best combination of sensitivity and discriminative power was used for all subsequent protein- and peptide-level validation.

Having identified *DWrollup* as the most effective protein summarization strategy, its quantitative accuracy and specificity were evaluated at both the protein and peptide levels. At the protein level, 278 *E. coli* spike-in proteins and 1,567 human background proteins passed filtering (1,845/2,562 total proteins retained). QUANTIX recovered a mean observed log_2_ fold-change of 1.069 relative to the expected value of 1.0 (mean error = +0.069, RMSE = 0.576; *p* = 0.045, one-sample *t*-test against the expected fold-change), and achieved an AUC of 0.891 for distinguishing spike-in from background proteins (Fig. 6a,c). Direction-aware sensitivity was 50.7%, with 141 of 278 *E. coli* proteins detected at Benjamini-Hochberg (BH) FDR-adjusted *q <* 0.05, while the false-positive rate among human background proteins was 2.68% (42 of 1,567 proteins).

**Fig. 6.**
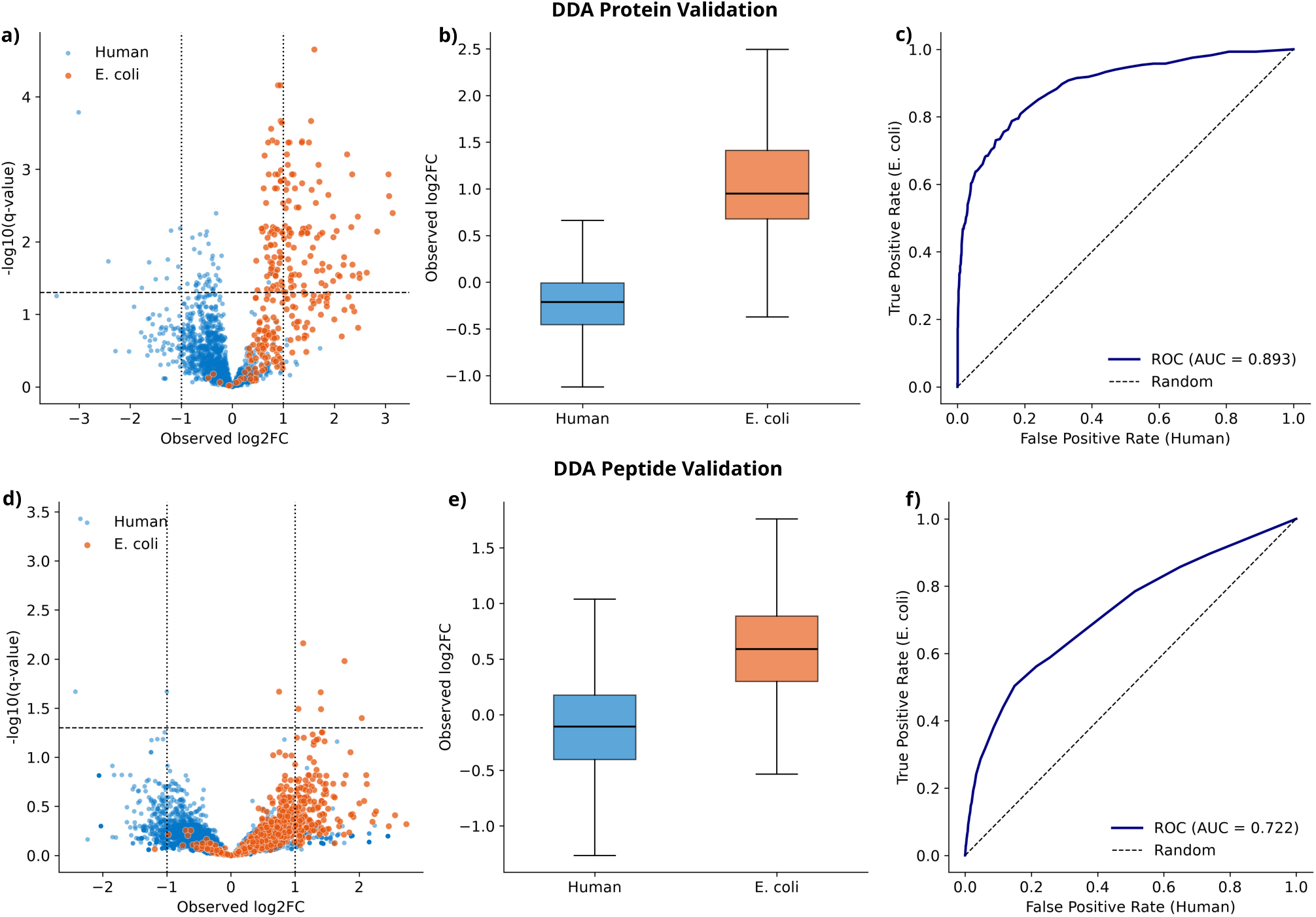
QUANTIX quantification is accurate and specific on a ground-truth benchmark. **a–c** Protein-level validation on the *E. coli* spike-in benchmark (2× versus 1× spike-in into a constant HEK293 background, PXD018408). (**a**) Volcano plot of observed log_2_ fold change versus − log_10_(*q*-value) for human (blue) and *E. coli* (orange) proteins. The dashed horizontal line indicates the significance threshold, and the dotted vertical lines indicate the ±1 log_2_ fold-change threshold. *E. coli* proteins, expected to exhibit a true twofold change, are enriched at positive fold changes relative to the human background, which is expected to show no change. (**b**) Distribution of observed log_2_ fold changes for human and *E. coli* proteins. Boxplots show the median, interquartile range, and 1.5× IQR whiskers, demonstrating separation between the expected-null human population and the expected twofold-change *E. coli* population. (**c**) Receiver operating characteristic curve for discriminating *E. coli* from human proteins by observed fold change (AUC = 0.893). (**d–f**) Peptide-level validation. (**d**) Volcano plot. (**e**) Distribution of observed log_2_ fold changes by species. (**f**) ROC curve (AUC = 0.722). Compared with protein-level quantification (**a–c**), peptide-level measurements show weaker, but still above-chance, discrimination between *E. coli* spike-in and human background peptides. Quantification was performed using the *DWrollup* strategy.

At the peptide level, 904 *E. coli* spike-in peptides and 5,959 human background peptides passed filtering (6,863/25,833 total peptides retained). Peptide-level quantification showed substantially reduced accuracy relative to the protein level, with a mean observed log_2_FC of 0.611 (mean error = −0.390, RMSE = 0.622; *p <* 0.001), reflecting systematic underestimation of the true fold-change when aggregating at the peptide rather than protein level (Fig. 6d,f). Discriminative performance likewise decreased, with an AUC of 0.722 and a sensitivity of only 1.1% (10/904 *E. coli* peptides at adjusted *q <* 0.05), though the false-positive rate remained very low (0.07%, 4/5,959 human peptides).

## 3 Discussion

MSGFLEX is an end-to-end proteomics workflow that integrates database searching (SPARX), machine learning-based PSM rescoring (XPECTRA), and protein quantification (QUANTIX) within a single deployable framework. Unlike conventional workflows, in which individual tools are assembled by users across disparate file formats[20–22], MSGFLEX maintains a common representation of identifications throughout the pipeline, enabling direct assessment of how changes in PSM confidence propagate to downstream protein-level quantification — an evaluation that is particularly relevant in metaproteomics, where database complexity, identification confidence, and quantitative inference are closely associated.

Building on this integrated framework, XPECTRA was benchmarked against Percolator, MS^2^Rescore, and MSBooster+Percolator across datasets spanning synthetic communities, human gut microbiomes, marine metaproteomes, and human proteomes — contexts in which the benefits of rescoring have been previously documented[3, 7, 8, 23, 24]. Across all datasets, XPECTRA consistently produced the largest identification gains and effect sizes among the methods evaluated, recovering 231,696 additional target PSMs on the human gut metaproteome dataset (PXD035981) alone. The consistency of these gains across datasets with substantially different sample compositions and proteome complexity suggests that XPECTRA captures discriminative information that generalizes beyond any particular biological system, though the magnitude of improvement remained dependent on the characteristics of the underlying spectra and search results. XPECTRA was developed and benchmarked exclusively on DDA data; extending the framework to data-independent acquisition workflows or alternative search engine output formats represents an important direction for future work.

Recent rescoring frameworks, including MSBooster and MS^2^Rescore, increasingly depend on external peptide-property predictors — such as DeepLC, MS^2^PIP, Prosit, or Koina-hosted models — to generate fragment-ion intensity and RT predictions incorporated into rescoring [25–28]. XPECTRA instead derives all features directly from database-search results and experimentally observed spectra, using XGBoost as the primary discriminative model to integrate complementary spectral, chromatographic, and precursor-derived evidence without reliance on external prediction infrastructure. The performance of XPECTRA across diverse datasets suggests that classical machine-learning frameworks remain competitive with deep-learning-augmented rescoring approaches when informative experimental and search-derived features are available[4], a consideration particularly relevant for large-scale metapro-teomic searches where pipeline simplicity and reproducibility are prioritized.

The consistency of SHAP-based feature importance across datasets supports the interpretation that XPECTRA learned generalizable relationships between spectral quality and identification confidence, rather than overfitting to dataset-specific score distributions. This is particularly relevant for metaproteomics, where training data characteristics can vary substantially across sample types and search databases. The divergence between SHAP and gain-based rankings for features such as RT deviation further illustrates why interpretability tools beyond split-based importance are valuable for understanding machine-learning rescoring models[13]. The noisy-OR fusion step introduces a layer between constituent model predictions and the final confidence score, limiting direct feature attribution at the fused-score level; SHAP analysis of the pre-fusion ensemble partially addresses this, and future designs could aim for more transparent attribution through the full scoring procedure.

XPECTRA incorporates several design features that distinguish it from existing rescoring approaches. Unlike Percolator, MS^2^Rescore, and MSBooster, which evaluate each PSM independently, XPECTRA includes leakage-controlled, trainingfold-derived peptide statistics that capture the consistency of high-confidence peptide observations across the dataset. Spectral features are derived directly from experimentally observed spectra and established fragmentation principles, including the proline effect[29] and mobile-proton-model-dependent cleavage enhancement at acidic residues under low-charge conditions[30]. Finally, ensemble-derived probabilities are combined with native MS-GF+ spectral evidence through noisy-OR fusion, ensuring that search-engine evidence contributes directly to the final score alongside machine-learning-based discrimination. Together, these choices allow XPECTRA to integrate cohort-level, spectral, and search-engine-derived evidence within a single rescoring framework.

Large multi-organism sequence databases represent a key computational bottleneck in metaproteomics, as searches against millions of protein sequences impose substantial memory and runtime demands[31, 32]. SPARX addresses this through database binning and resource-aware search scheduling, partitioning large databases into independent subsets while controlling computational resource requirements. Critically, concordance between binned and unified database searches was evaluated after XPECTRA rescoring — the workflow configuration used for downstream analyses — rather than at the native MS-GF+ stage alone. The high concordance in score distributions, peptide assignments, and target-decoy confidence estimates under these conditions indicates that database binning can improve scalability without introducing detectable bias in rescoring performance. Whether this behavior generalizes to substantially larger, more redundant, or taxonomically broader databases will require further evaluation.

Benchmarking against the ground-truth spike-in dataset (PXD018408) indicated that *DWRollup* provided the most favorable balance of sensitivity, specificity, and discriminative performance among the evaluated protein rollup strategies. To our knowledge, this represents the first empirical validation of a degeneracy-weighted rollup approach for intensity-based protein quantification using known spike-in standards. Comparisons with alternative methods, including distributed Normalized Spectral Abundance Factor (dNSAF)[33] and the peptide-scaling approaches *Rrollup*, *Zrollup*, and *Qrollup*, further demonstrated that quantitative outcomes can be strongly influenced by implementation details in addition to the underlying methodology. In particular, *Zrollup* underperformed because normalization was applied along an unsuitable axis, reducing biologically meaningful fold-change information, whereas *Qrollup* differed substantially from the original DAnTE implementation because standard quantiletransformation routines do not faithfully reproduce the published algorithm[34]. These observations highlight a broader challenge in quantitative proteomics: methods sharing the same name can produce substantially different results when implementation choices, normalization procedures, or statistical assumptions vary[35]. Together, these findings underscore the importance of benchmarking quantitative workflows against ground-truth datasets and suggest that *DWRollup* represents a promising approach for protein-level quantification. Nevertheless, validation was performed using a single spike-in dataset (PXD018408), and evaluation across additional benchmark datasets spanning different background complexities and spike-in ratios will be important for assessing its broader generalizability.

Taken together, these results establish MSGFLEX as a unified framework in which database searching, PSM rescoring, and quantitative analysis can be evaluated both independently and as an integrated workflow. Rather than validating each component in isolation, interactions between workflow stages were examined directly, including the effects of XPECTRA rescoring on partitioned database searches and the impact of QUANTIX quantification on rescoring-filtered identifications. This distinction is important because gains in PSM identification do not necessarily translate into improved protein-level quantification if additional identifications are unevenly distributed across proteins or introduce quantitative uncertainty. By evaluating these dependencies within a common analytical framework, MSGFLEX provides a more realistic assessment of practical performance than component-level benchmarking alone. Furthermore, its modular design enables independent extension of individual components, such as incorporating additional SPARX-derived features into XPECTRA or implementing new protein rollup strategies in QUANTIX, providing a flexible foundation for future developments in metaproteomics and microbiome research.

**Deployability.** MSGFLEX was designed with practical throughput and computational scalability in mind. In particular, its NUMA-aware scheduling strategy assigns MS-GF+ search jobs according to the available NUMA topology, matching thread and task allocation to the detected hardware configuration. By determining the system topology at runtime through _get_core_config(), SPARX can adapt its parallelization strategy to multi-socket systems without requiring users to manually configure hardware-specific settings. This design reduces the need for manual resource tuning and provides a more reproducible approach to deploying large database searches across heterogeneous computing environments. Implementation details of the scheduling strategy are provided in the Supplementary Methods. Currently, MSGFLEX supports Thermo raw files and has been validated on Linux; extension to additional raw file formats and UNIX-like platforms remains ongoing.

## Acknowledgments

This work was supported by the National Institutes of Health award number is R01AT013241. The authors also acknowledge the Information Technology Services at Baylor University for providing High-Performance and Research Computing Services (Kodiak), which have contributed to the research results reported within this paper; URL: https://its.web.baylor.edu/hpc.

## Author contributions

A.T.W convceived the conceptualization. T.R.R, H.A.H, and A.T.W. designed the methodology. T.R.R. and H.A.H. wrote the code and performed the software imple-mentation. T.R.R. designed the experiments and performed the formal analysis and validation. A.T.W. supervised the project. T.R.R. wrote the original draft of the manuscript. A.T.W, H.A.H, and T.R.R. authors reviewed and edited the manuscript.

## Data availability

The datasets analyzed in this study are publicly available from the PRIDE repository and MassIVE under accession numbers PXD006118, PXD017035, PXD018408 (MSV000085239), PXD035981, PXD057701, and PXD074461. Proteomics data generated with the MSGFLEX pipeline have been deposited to the ProteomeXchange Consortium via the PRIDE repository[36] and MassIVE; accession numbers and repository details are provided in Supplementary Table S3. These datasets are currently private and will be made publicly available upon acceptance; reviewer access tokens are provided in Supplementary Table S3. Additional scripts and metadata supporting the findings are available at https://github.com/WrightLab-Baylor/msgflex_manuscript.

## Code availability

MSGFLEX, including the SPARX, XPECTRA, and QUANTIX modules, is implemented in Python and is publicly available at https://github.com/WrightLab-Baylor/msgflex. The version used for all analyses reported in this study is archived as release v1.1.0. Installation instructions, usage examples, and detailed documentation are provided in the repository and Supplementary Methods. An optional desktop interface based on DearPyGui is also provided. MSGFLEX integrates several third-party tools, including ThermoRawFileParser, MS-GF+, PHRP, and MASIC, which must be obtained separately under their respective licenses. MSGFLEX is distributed under the Apache License 2.0.

## 4 Methods

MSGFLEX is an end-to-end computational framework for metaproteomic analysis comprising three modules: SPARX for raw-file processing and MS-GF+ database searching, XPECTRA for machine-learning-based PSM rescoring, and QUANTIX for peptide- and protein-level relative quantification (Fig. 1). SPARX generates feature-rich MS-GF+ PSM tables and chromatographic measurements that serve as input to XPECTRA. XPECTRA integrates search-engine, spectral, chromatographic, and peptide-level evidence to rescore PSMs using a leakage-controlled ensemble learning framework. QUANTIX uses the resulting peptide identifications and chromatographic intensities for downstream peptide- and protein-level quantification.

### 4.1 SPARX: raw file conversion and database search

The SPARX module (designated as STAGE 1 in Fig. 1) performs upstream spectral processing and database searching, converting vendor-specific raw spectral files into feature rich peptide-spectrum match (PSM) tables ready for downstream analysis, including rescoring. The SPARX module is designed exclusively for Linux-based systems, with proprietary Windows-native tools (e.g., ThermoRawFileParser, PHRP) invoked through the Mono runtime.

Vendor-specific Thermo RAW files are converted to mzML format using ThermoRaw-FileParser [37], under the Mono runtime. The resulting mzML files are input for MS-GF+ database searching. The resulting mzML files are parsed with Pytoemics package to generate QC plots[38].

SPARX supports two database-search modes. In conventional mode, MS-GF+ searches the complete protein FASTA database in a single pass. In binning mode, the database is partitioned into smaller sub-databases searched independently and subsequently aggregated. Database binning is designed for large metaproteomic sequence databases implemented primarily to address a well-documented sensitivity limitation of standard database search engines including MS-GF+, when applied to very large sequence databases (>10^6^ entries) typical of multi-organism metaproteomics datasets[5, 6]; partitioing the search space reduces the scoring burden and improves sensitivity with peptide recovery relative to a single-pass search.

SPARX performs database searching using MS-GF+, chosen for its high sensitivity, robust scoring model, and generating-function-based statistical framework, which produces calibration-free spectral significance estimates[39, 40]. These properties make MS-GF+ particularly well suited for the large, heterogeneous, and often taxonomically uncharacterized sequence databases commonly encountered in metaproteomics. Target-decoy search was enabled (-tda 1) to facilitate downstream FDR estimation, and the –addFeatures flag was also enabled to generate a feature-rich PSM table for rescoring. Search parameters including precursor mass tolerance, enzyme specificity, and fixed/variable modifications explicitly configurable via designated MS-GF+ parameters file (e.g., params.txt). MS-GF+ mzIdentML (.mzid) output was converted to tab-separated format (.tsv) using MS-GF+’s native conversion utility.

Raw MS-GF+ output reports PSMs strictly at the spectrum level, with no explicit unique peptideor protein-level aggregation, nor it does account for global false discovery rate (FDR) across complex multi-sample datasets. To standardize search-engine output and facilitate downstream rescoring and quantification, SPARX invokes the Peptide Hit Results Processor (PHRP), which resolves peptide-to-protein map-pings, applies mass corrections, and recalculates *q*-values. [https://github.com/PNNL-Comp-Mass-Spec/PHRP].

Selected-ion chromatogram (SIC) statistics are necessary for downstream rescoring and peptide-to-protein relative quantification. SPARX module indepently extracsts SIC features from each raw file with MASIC [41]. These measurements included precursor- and selected-ion-chromatogram statistics used by both XPECTRA rescoring and QUANTIX quantification. A custom MASIC Merger component integrated MASIC-derived features with the corresponding PHRP-processed PSM tables.

SPARX is designed to be run on proteomic high-end workstations with minimum of 64GB RAM and 16 CPU cores, or high-performance computing (HPC) clusters.

SPARX supports parallel execution of database searches across available compute resources; implementation details and hardware-specific optimization are described in Supplementary Methods.

*Dynamic Resource Scheduling.* To optimize the MS-GF+ database search speed and to utilize the available computing resources efficiently, SPARX performs runtime resource-aware scheduling. At execution, the pipeline detects available system resources, including physical core count, NUMA topology, and total system memory. Thread and task allocations are distributed across concurrently executing searches such that aggregate thread demand does not exceed the available physical cores. The maximum number of simultaneous jobs is additionally constrained by available memory, calculated as 90% of total RAM divided by the configured Java heap size per search process.

### 4.2 XPECTRA: cross-validated ensemble rescoring

XPECTRA is a machine-learning-based PSM rescoring module that integrates complementary evidence from MS-GF+ search scores, tandem-mass-spectral features, RT information, and peptide- and precursor-level characteristics. It was implemented in Python using XGBoost as the core classification model; the complete software environment and package versions are available in the project’s GitHub repository (Code Availability). PSMs are rescored using a leakage-controlled, cross-validated ensemble of gradient-boosted decision-tree classifiers, producing held-out predictions for every PSM. These predictions are subsequently combined with native MS-GF+ spectral evidence to generate the final XPECTRA score used for FDR-controlled identification. The detailed XPECTRA rescoring architecture is illustrated in Fig. 7. XPECTRA recognizes modified peptides directly from the modification annotations (mass shifts) reported in the PHRP-processed search results, so no external configuration or user-supplied modification table is required. These annotations are used to correct predicted RTs for common modifications that alter peptide elution: methionine oxidation, phosphorylation, deamidation, protein N-terminal acetylation, and lysine acetylation that blocks TMT labeling. Because modification information is read from the search output itself, the same model applies to unlabeled and TMT-labeled datasets without dataset-specific feature engineering.

**Fig. 7.**
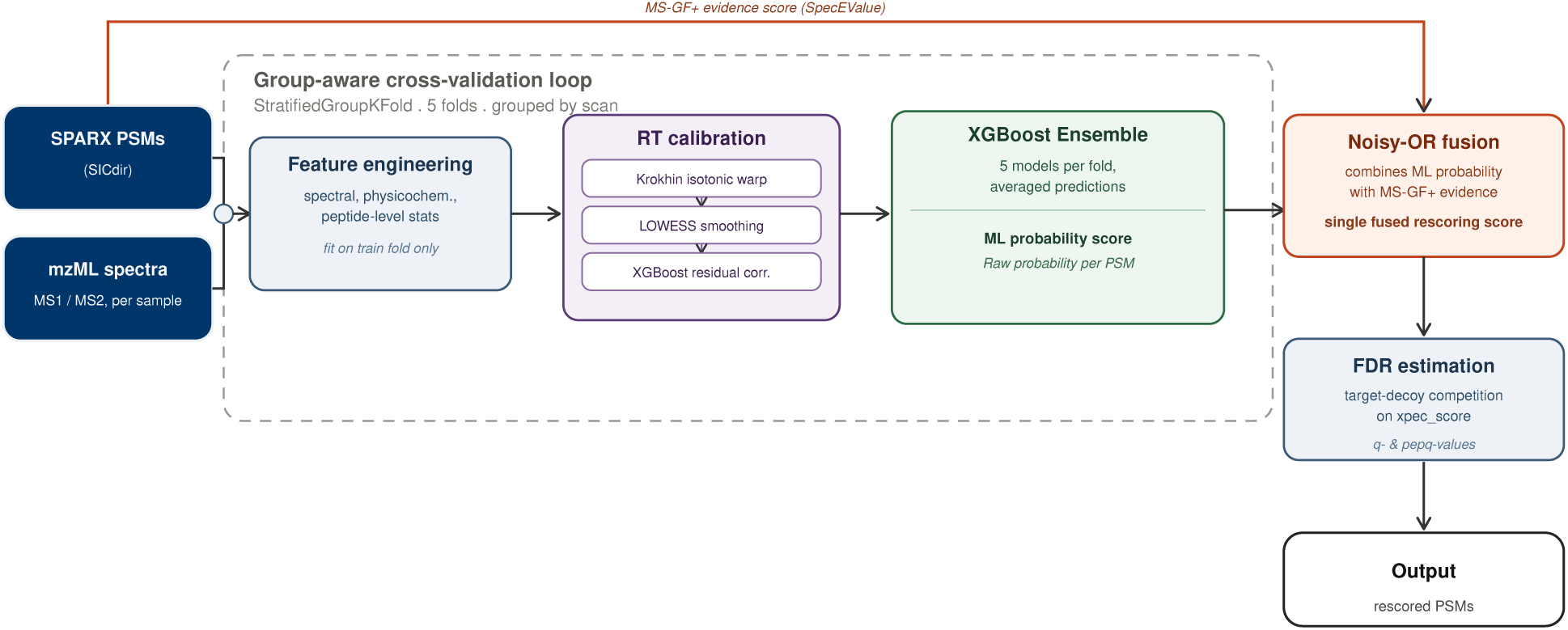
Detailed XPECTRA rescoring architecture. SPARX-generated PSMs (with associated selected-ion chromatogram directory, SICdir) and raw mzML spectra (MS1/MS2, per sample) feed into a group-aware cross-validation loop using StratifiedGroupKFold (5 folds, grouped by scan to prevent data leakage between folds). Within each fold, features are engineered from spectral, physicochemical, and peptide-level statistics, fit on the training fold only; retention time is calibrated via Krokhin isotonic warping[47] with LOWESS smoothing and XGBoost residual correction. A five-model XGBoost ensemble per fold produces averaged raw probability predictions per PSM. These ML probabilities are combined with MS-GF+ evidence scores (SpecEValue) via noisy-OR fusion to yield a single fused rescoring score (xpec_score). Target-decoy competition on this fused score produces final *q*-values and peptide-level *q*-values (pepq), yielding the final set of rescored PSMs.

For each PSM, XPECTRA constructs a feature vector comprising four categories of evidence: (i) native MS-GF+ search scores, including transformed spectral scores and E-values; (ii) tandem-mass-spectral evidence derived from matched theoretical and observed fragment ions; (iii) RT features based on predicted-versus-observed elution behavior; and (iv) peptide- and precursor-level characteristics, including precursor mass accuracy, peptide properties, and peptide observation statistics. During feature extraction, candidate *b*- and *y*-ion fragments are additionally weighted using established CID fragmentation principles, including the proline effect and mobileproton-model-dependent cleavage enhancement at acidic residues under low-charge conditions[29, 30]. XPECTRA also incorporates Kendrick mass defect (KMD) variance as an additional spectral feature[42], and identify identify atypical isotope distribution patterns of precursor ions with averagine approximation[43]. To avoid train — test leakage, feature transformations and any empirically derived parameters were fit using only the training data within each cross-validation fold and subsequently transferred unchanged to the held-out fold during inference.

RT deviation is used as an additional PSM feature. Peptide RTs were first predicted from sequence-derived hydrophobicity and subsequently calibrated using training-fold data to account for systematic RT behavior and relevant peptide properties. The absolute deviation between predicted and observed RT was then calculated and incorporated into the XPECTRA feature vector. Details of the RT prediction and calibration procedure are provided in Supplementary Methods.

To prevent spectrum-level information leakage, PSMs were partitioned into five group-stratified folds using OptimalScanNumber as the grouping variable and target/decoy status for stratification. For each fold, five independently seeded XGBoost models were trained on the remaining data and used to score the test fold. Predictions were aggregated using a trimmed mean to produce the final out-of-fold rescoring probability (rescore_prob) for each PSM.

The final XPECTRA score is denoted as xpec_score, an ensemble-derived score was combined with calibrated MS-GF+ spectral evidence using a noisy-OR fusion function[15]. Let *p* denote the ensemble-derived confidence socre and *e* the calibrated native MS-GF+ spectral evidence. The resulting xpec_score is calculated as Eq.(2):

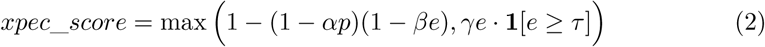

The parameters *α* and *β* control the relative contribution of the learned and native evidence, respectively. A conditional rescue term was included to retain PSMs with strong native MS-GF+ evidence despite a low ensemble score: *τ* is the spectral-evidence threshold above which the rescue term activates, and *γ* is the rescue coefficient, determining what fraction of spectral evidence *e* is guaranteed to the final score once *e ≥ τ* (*γ* = 0.80, *τ* = 0.70 used throughout)

PSM- and peptide-level *q*-values are calculated from the fused score using standard target-decoy competition [44], following the implementation used by Feng et al.[32]. False discovery rates are estimated as in Eq. (3):

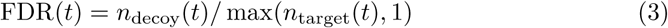

and monotonic *q*-values are obtained using a standard tail-minimum correction. Peptide-level *q*-value are computed using the highest-scoring PSM for each peptide and then assigned back to all corresponding PSMs.

*Evidence-weight selection.* The evidence weight selected independently for each dataset via grid search over *α ∈ {*0.25, 0.5, 0.75, 1.0, 1.25, 1.5, 2.0*}*, while *β* was held fixed at 1.0. For each candidate value of *α*, the fused score is computed for all PSMs, PSM-level-*q*-values are estimated by target-decoy competition, and the number of target PSMs accepted at 1% FDR is recorded. The value of *α* yielding the highest number of accepted target PSMs is selected for final scoring. This dataset-specific optimization allows the contribution of ensemble-derived and native spectral evidence to be adjusted according to the quality of the underlying scores in each dataset.

*Design philosophy and interoperability.* XPECTRA was designed as a modular upstream rescoring engine that terminates at the peptide level. Protein inference was intentionally excluded from the rescoring stage to avoid imposing assumptions regarding shared peptides, protein isoforms, and protein grouping strategies. By separating spectral confidence estimation from protein inference, XPECTRA can be coupled with a variety of downstream protein-level analysis frameworks. Accordingly, XPECTRA outputs standard tab-delimited files that can be used directly by established protein inference tools and quantification workflows.

### 4.3 QUANTIX: peptide-to-protein relative quantification

The third module of MSGFLEX framework is QUANTIX, which consolidates peptide and protein abundances from SPARX or XPECTRA output. Parent-ion and selected-ion-chromatogram (SIC) intensity measurements extracted by MASIC are used as the basis for peptide-level quantification. SPARX uses PHRP generated first-hit-table _fht.txt as the input for MASIC merging, wheras QUANTIX additionally incorporates peptide assignments from the corresponding PHRP generated synopsis files _syn.txt. The _syn records provide peptide-level associations for co-eluting peptide species, including peptides that were not directly selected for MS/MS but share chromatogrphic evidence with confidently identified peptides. QUANTIX merges these peptide assignments with MASIC-derived SIC measurements to construct the peptide-by-sample abundance matrix.

PSMs are filtered before quantification using a user-specified confidence metric and threshold. Supported filtering metrics include MS-GF+’s native spectral E-value, exposed within the pipeline as MSMSScore (− log_10_(SpecEValue), such that higher values indicate greater confidence), as well as PSM-and peptide-level *q*-values and XPECTRA-derived scores (xpec_q, xpec_pepq, or xpec_score).

Following confidence PSM filtering, peptide intensities from all samples are combined into a single peptide-by-sample abundance matrix. Each peptide is then mapped to the reference protein database and annotated with the corresponding protein accession, gene name, and functional description. Peptides are classified as either unique or shared based on their protein assignments: peptides mapping to a single protein are considered unique, whereas those mapping to multiple proteins are considered shared.

QUANTIX implements the three protein rollup strategies originally described in DAnTE (*RRollup*, *QRollup*, and *ZRollup*) and subsequently adopted in pmartR[34, 45]. In addition to these methods, QUANTIX supports direct intensity summation (*Sum*) and a new Degeneracy-Weighted Rollup (*DWRollup*) strategy. Peptide *p*’s intensity is denoted *x_p,s_* on the log_2_ scale and *I_p,s_* = 2*^xp,s^* on the linear scale, and *P_g_* denotes the set of peptides mapped to protein *g*.

In the *Sum* method, the linear-scale intensities of all peptides assigned to a protein are summed directly:

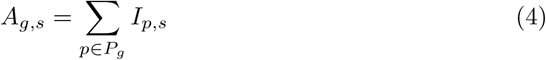

*Rrollup* selects a reference peptide *r* for each protein, chosen as the peptide with the fewest missing values (or, in the case of a tie, the highest median intensity across samples). Over the set of samples 0*_p_* in which both peptide *g* and the reference *r* are observed, the reference-scaled intensity *x_p,s_* is computed by shifting *g*’s log_2_ intensities to the reference peptide’s level, and the protein abundance is the median of the scaled peptide intensities as Eq. (5):

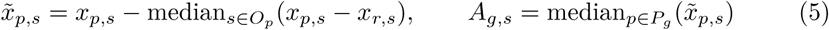

An optional Grubbs’ test can be applied to exclude outlier values before the median calculation [34, 46].

*Zrollup*, inspired by DAnTE[34] but implemented here using mean/SD standardization rather than DAnTE’s median-based scaling, converts each peptide’s intensities to a standardized (z-scored) intensity *z_p,s_* and takes the median z-score per protein:

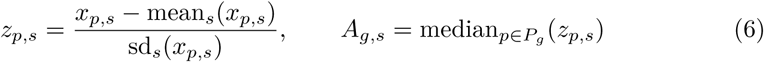

*Qrollup* ranks peptides by their mean intensity across samples and retains, in the set *Q*, those peptides whose mean intensity meets or exceeds a threshold *τ* set to the 1 − *f*-th percentile of peptide means (*f* = 1/3 by default, *i.e.* the top third of peptides by mean intensity); the protein abundance is the mean of the retained peptides’ intensities in each sample, reducing the influence of low-abundance peptides on the protein-level estimate[34]:

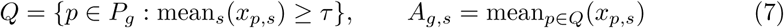

We additionally introduce a fifth protein rollup strategy, *DWRollup*, which explicitly accounts for peptide-sharing ambiguity. Peptide-protein degeneracy is computed across the complete annotated peptide-protein graph, and each peptide is assigned a weight inversely proportional to *n_p_*, the number of distinct proteins to which peptide *p* maps. Weights are subsequently normalized within each protein such that ∑_pɛPg_*w_p_* = 1, and protein abundance is calculated as the weighted average of constituent peptide intensities. Consequently, peptides shared across multiple proteins contribute proportionally less to protein abundance estimates than peptides uniquely assigned to a single protein, while still retaining quantitative information as in Eq. (8).

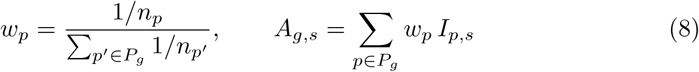

All five methods can be run in one of three modes: keep all peptide-protein matches regardless of sharing (*all_matches*), use only unique peptides (*unique_only*), or only quantify a protein if every one of its peptides is unique to it (*requires_unique*).

QUANTIX does not perform cross-sample normalization or missing-value imputation. The resulting peptide- or protein-level abundance matrices preserve the measured intensity values and can therefore be subjected to downstream normalization, imputation, and statistical analysis according to the experimental design.

### 4.4 SHAP-based feature attribution

To interpret what drove XPECTRA’s rescoring decisions, we computed SHAP values[13] for every PSM, using the tree-based explainer (TreeExplainer)[14], which is exact for gradient-boosted tree models like XGBoost rather than relying on an approximation.

Because each ensemble fold consisted of five independently seeded XGBoost models, SHAP values were calculated separately for each model using PSMs from the corresponding held-out fold. Feature attributions were then averaged across the five ensemble members and assigned to each PSM. This approach replicates the generation final rescoring probability score, which is similarly derived from an ensemble average, ensuring that the reported SHAP values explain the score ultimately used for rescoring rather than the predictions of any single model.

Then, SHAP values were then analyzed alongside the corresponding input features utilized during prediction to assess both the magnitude and direction of their impact, revealing whether high or low feature values increased or decreased the likelihood of PSM acceptance.

### 4.5 Benchmarking against Percolator, MS^2^Rescore, and MSBooster

XPECTRA’s rescoring performance was compared against three widely used PSM rescoring tools - Percolator[2, 3], MS^2^Rescore[7], MSBooster[8] with Percolator - alonside the unrescored MS-GF+ baseline (before PHRP normalization) and un-rescored SPARX baseline (Post PHRP normalization). All four rescoring models recieved the same underlying MS-GF+ PSMs as input for each sample, ensuring that observed differences in identification counts arose from the rescoring model implementation rather than underlying search results.

XPECTRA of MSGFLEX follows non-parsimoniuos protein inference while Percolator or Mokapot of MS^2^Rescore follows parsimonious protein inference. As a result, identical peptide sequences may be assigned to different proteins under parsimonious inference, depending on the inferred minimal protein set. To elimate the mapping artifcat, a single consensus peptide-to-protein map was constructed prior to comparison: for every peptide observed across all MS-GF+ output files, its most frequently co-occuring protein accession (by majority vote across samples) was designated canonical.

Rescoring performance was evaluated at multiple FDR thresholds (1%, 2.5%, and 5%) by comparing the numbers of accepted target PSMs, unique peptides, and unique consensus-mapped proteins obtained from each tool. Identification counts were determined using each tool’s native *q*-value estimates and target-decoy classification. For MS^2^Rescore, which does not provide peptide-level *q*-values, peptide-level *q*-values were computed by selecting the highest-scoring PSM for each peptide and applying standard target-decoy competition, consistent with the peptide-level FDR estimation procedure used for XPECTRA. Counts were calculated on a per-sample basis and summed across samples to generate experiment-level totals. Relative to the MS-GF+ baseline, accepted identifications from each rescoring tool were further classified as gained, lost, or shared. Gain/loss analyses were performed using set-based comparisons of accepted target PSMs, peptides, and proteins at each FDR threshold, matching identifications by spectrum scan number whenever correspondence could be established across tool output formats and by aggregate identification counts otherwise. Exact version numbers for all third-party tools used throughout MSGFLEX are listed in Table S2, including MS-GF+, ThermoRawFileParser, PHRP, MASIC, Percolator, MS^2^Rescore, and MSBooster.

