## Supplementary Information for "MSGFLEX: An optimized end-to-end pipeline for proteomics and metaproteomics analysis built on MS-GF+ with a dedicated rescoring model, XPECTRA"

**Table S1. Software and package versions used in this study.**

| Software/Package | Version | Purpose | Reference |
|---|---|---|---|
| *Raw data processing & database search* | | | |
| pyteomics | 4.7.5 | Parse mzML files | Goloborodko et al., 2013 |
| ThermoRawFileParser | 1.4.2 | Raw file conversion | Hulstaert et al., 2019 |
| MS-GF+ | 2024.03.26 | Search Engine | Kim and Pevzner, 2014 |
| PHRP | 3.1.8056.28963 | MS-GF+ Peptide Hit Process | -NA- |
| MASIC | 3.2.8152.28867 | SIC statistics | Monroe et al., 2008 |
| *Benchmarking comparators* | | | |
| Percolator | 3.09.0 | PSM Rescoring | The et al., 2016 |
| MS$^2$Rescore | 3.2.1 | PSM Rescoring | Burr et al., 2024 |
| MSBooster | 1.3.31 | PSM Rescoring | Yang et al., 2023 |
| DeepLC | 3.1.13 | RT predictor for MS$^2$Rescore | Bouwmeester et al., 2021 |
| DIA-NN | 1.8 (beta) | RT predictor for MSBooster | Demichev et al., 2020 |

**Table S2. Datasets used in this study for benchmark analysis.**

| PRIDE No. | Type | Samples used | Instrument | Modifications |
|---|---|---|---|---|
| PXD006118 | Synthetic community | 144 | Q Exactive | Acetylation (Protein N-term, K)<br>Oxidation (M)<br>Carbamidomethylation (C) |
| PXD017035 | SIHUMIx | 16 | Q Exactive | Acetylation (Protein N-term, K)<br>Oxidation (M)<br>Carbamidomethylation (C) |
| PXD018408 | HEK293 cells | 24 | Q Exactive | Acetylation (Protein N-term, K)<br>Oxidation (M)<br>Deamidation (N/Q)<br>Carbamidomethylation (C) |
| PXD035981 | Human gut microbiome | 24 | Q Exactive HF | Acetylation (Protein N-term, K)<br>Oxidation (M)<br>Deamidation (N/Q)<br>Carbamidomethylation (C) |
| PXD052873 | Ocean Metaproteome | 34 | Orbitrap Fusion Lumos | Acetylation (Protein N-term, K)<br>Oxidation (M)<br>Carbamidomethylation (C) |
| PXD057701 | Human gut microbiome | 8 | Q Exactive HF-X | Oxidation (M)<br>Carbamidomethylation (C) |
| PXD074461 | Human Proteome | 24 | Orbitrap Exploris 240 | Acetylation (Protein N-term, K)<br>Oxidation (M)<br>Carbamidomethylation (C) |

**Table S3. Accession details of the reanalyzed datasets.**

| Original Accession | Data Reference | Repository | Reanalysis Accession | Password/ Token |
|---|---|---|---|---|
| PXD006118 | Kleiner et al., 2017 | PRIDE | PXD085323 | 3kRMYOUDeUdW |
| PXD018408 | Dowell et al., 2021 | MassIVE | RMSV000000717 | WrightLabBU123 |
| PXD035981 | Raba et al., 2024 | PRIDE | PXD085450 | J7DkVQNkQIoD |
| PXD052873 | Mudge et al., 2025 | PRIDE | PXD085364 | gdTa0tqw86X0 |
| PXD057701 | Valdes-Mas et al., 2025 | PRIDE | PXD085497 | r1VHcEse4Qgq |
| PXD074461 | Costanzo et al., 2026 | PRIDE | PXD085424 | p6unmmtSgARG |

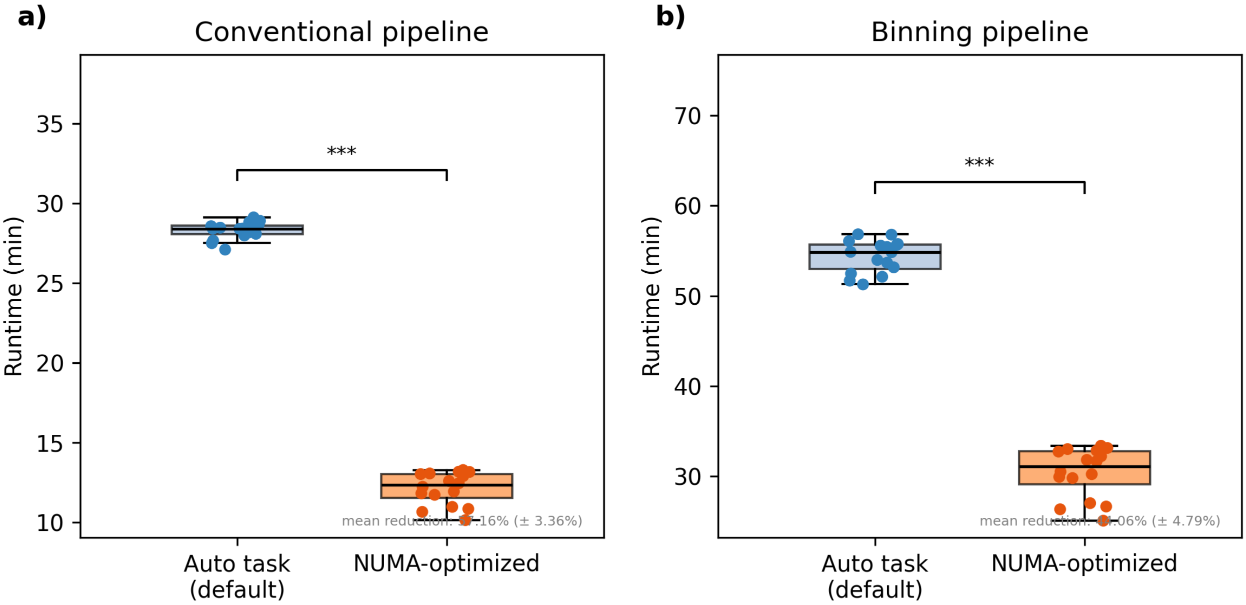


**Fig. S1.** Auto Task Optimization with NUMA-toplogy reduces the search time. **(a).** Runtime comparison between default task scheduling (Auto task) and NUMA-optimized core pinning for the conventional pipeline, benchmarked on [16] replicate runs per condition. NUMA optimization reduced mean runtime by 57.16% (± 3.36%). **(b).** Runtime comparison between default task scheduling (Auto task) and NUMA-optimized core pinning for the binning pipeline; NUMA optimization reduced mean runtime by 44.06% (± 4.79%). In both panels, boxes show the interquartile range with the median line, whiskers extend to the most extreme values within 1.5× IQR, and individual replicate runtimes are overlaid as points. Statistical significance was assessed by Wilcoxon signed-rank test; *** indicates $p <$ [0.001].

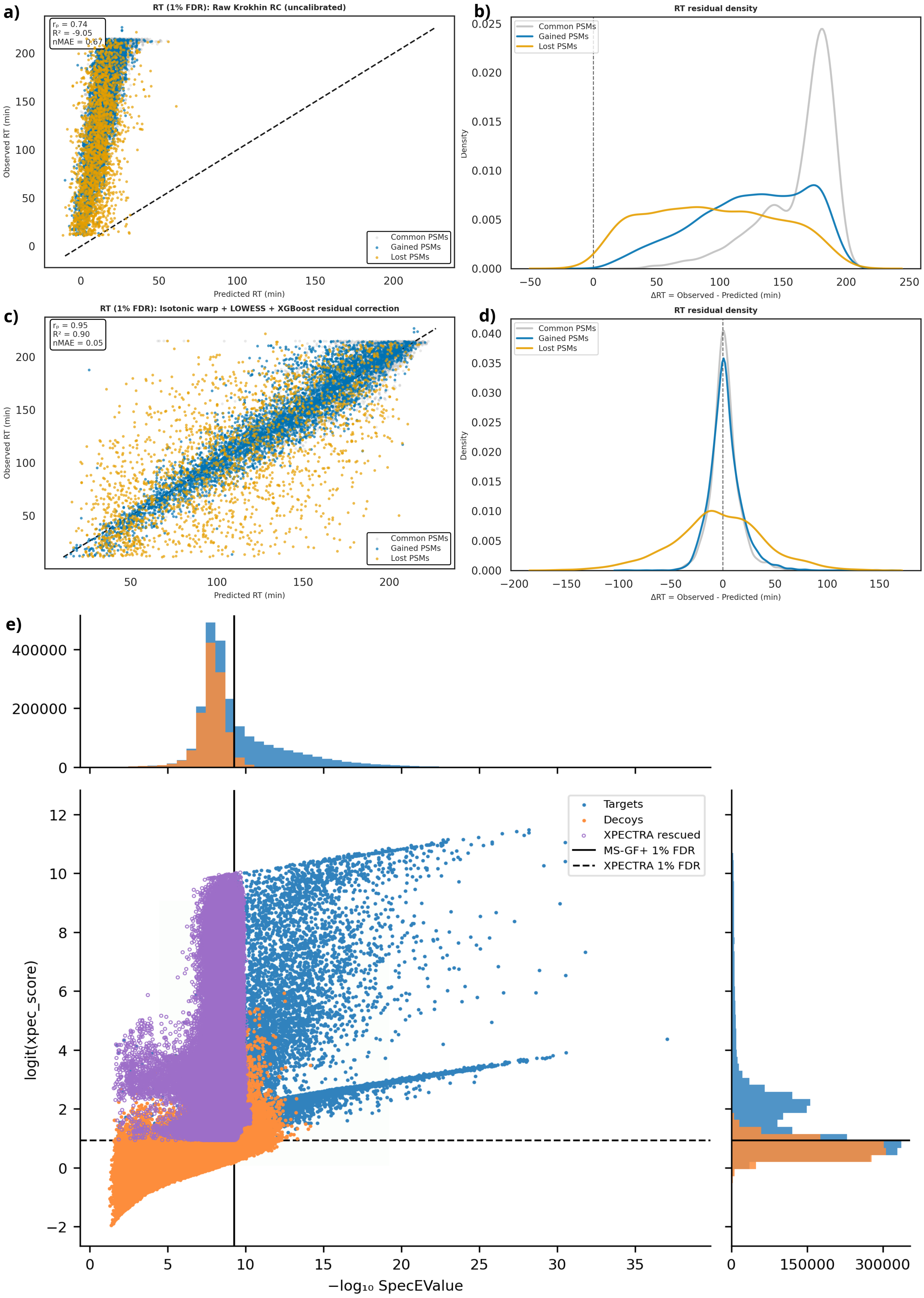


**Fig. S2.** Retention-time calibration and score distributions underlying XPECTRA rescoring. **(a)** Observed versus predicted retention time (RT) before calibration. Common, gained, and lost PSMs are shown in gray, blue, and orange, respectively; dashed line, $y = x$. **(b)** Distributions of RT residuals ($\Delta$RT = observed − predicted) before calibration. **(c)** Observed versus predicted RT after isotonic, LOWESS, and XGBoost-based calibration. Gained PSMs follow the diagonal as tightly as common PSMs, whereas lost PSMs are scattered widely off the diagonal. **(d)** RT residual distributions after calibration. **(e)** Score distributions for top-ranked PSMs of MS-GF+ SpecEValue ($-\log_{10}$ SpecEValue, x-axis) and XPECTRA rescoring score (logit(xpec_score), y-axis) for all target and decoy PSMs. Marginal histograms show the univariate distributions for each axis. Points are colored by category: targets (blue), decoys (orange), and PSMs rescued by XPECTRA (purple open circles) – target PSMs that fall below the MS-GF+ 1% FDR threshold (solid vertical line) but are recovered above the XPECTRA 1% FDR threshold (dashed horizontal line). The rescued population is concentrated at low SpecEValue confidence but high XPECTRA score, illustrating that XPECTRA identifies discriminative spectral and physicochemical signal not captured by the spectral E-value alone.

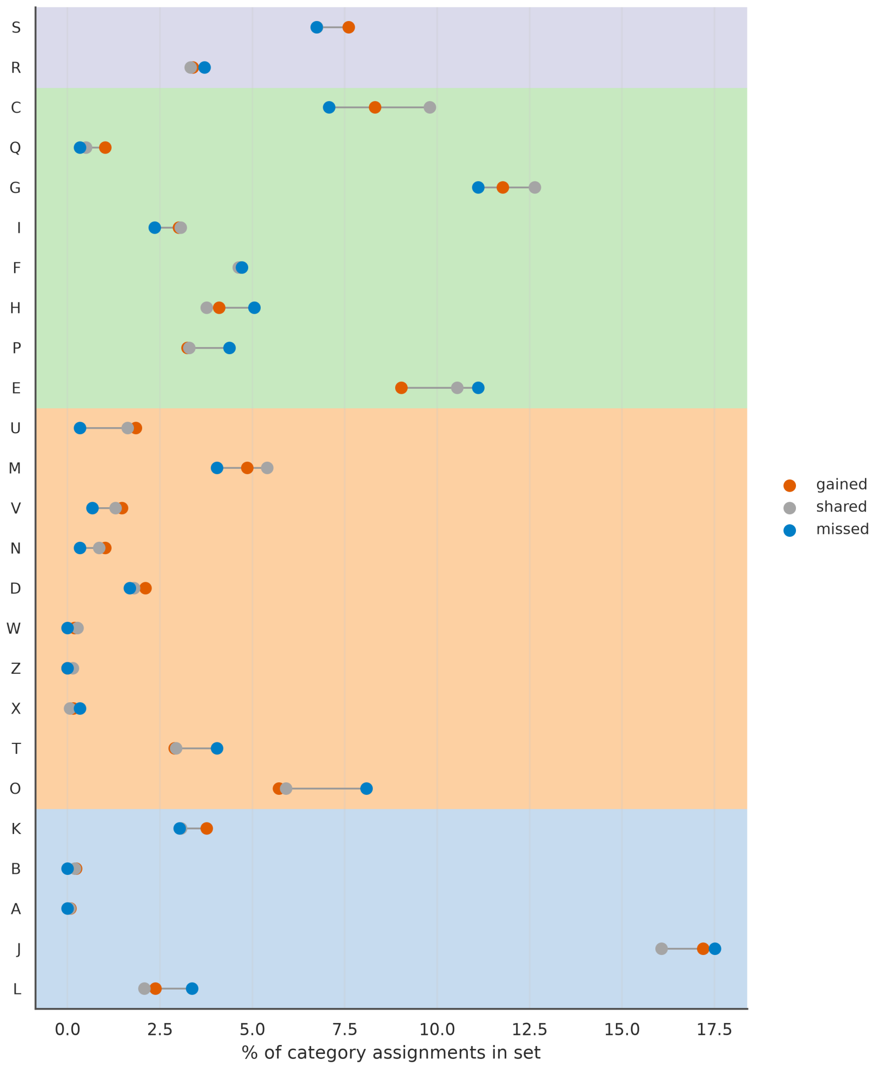


**Fig. S3.** Category-level distribution of proteins gained, shared, and missed by XPECTRA across COG functional categories. The Cleveland plot shows, for each COG category (rows), the percentage of total category assignments in the set attributable to XPECTRA gained (orange), shared (gray), and XPECTRA missed (blue) assignments. Categories are grouped into four functional supergroups, indicated by background shading: information storage and processing (light blue), cellular processes and signaling (light orange), metabolism (light green), and poorly characterized (light purple). The absence of strong category-specific enrichment suggests that XPECTRA gains are broadly distributed across functional classes rather than concentrated within specific biological processes. COG Category descriptions:
J – translation, ribosomal structure and biogenesis;
A – RNA processing and modification;
K – transcription;
L – replication, recombination and repair;
B – chromatin structure and dynamics;
D – cell cycle control, cell division, chromosome partitioning;
V – defense mechanisms;
T – signal transduction mechanisms;
M – cell wall/membrane/envelope biogenesis;
N – cell motility;
Z – cytoskeleton;
W – extracellular structures;
U – intracellular trafficking, secretion, and vesicular transport;
O – posttranslational modification, protein turnover, chaperones;
C – energy production and conversion;
G – carbohydrate transport and metabolism;
E – amino acid transport and metabolism;
F – nucleotide transport and metabolism;
H – coenzyme transport and metabolism;
I – lipid transport and metabolism;
P – inorganic ion transport and metabolism;
Q – secondary metabolites biosynthesis, transport and catabolism;
R – general function prediction only;
S – function unknown

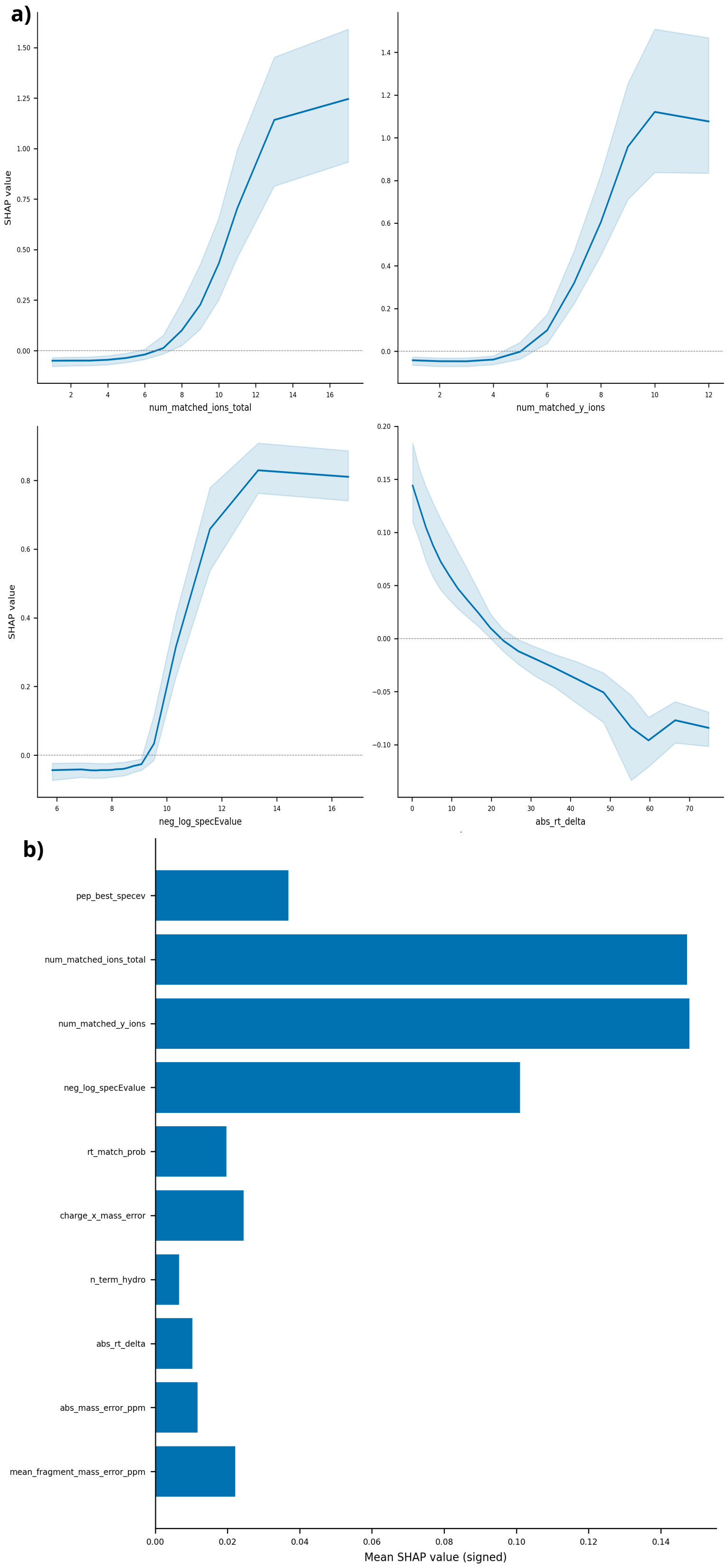


**Fig. S4.** XPECTRA feature-level SHAP behavior and XGBoost model importance ranking. **(a).** Mean SHAP value as a function of feature value (shaded band, [variability measure – e.g., ±1SD or 95% CI]) for the four top-ranked XPECTRA features: number of matched fragment ions (`num_matched_ions_total`), number of matched y-ions (`num_matched_y_ions`), MS-GF+ Spectral E-value ($-\log_{10}$ transformed; `neg_log_specEvalue`), and absolute retention time deviation (`abs_rt_delta`). The first three features show a positive, saturating relationship between feature value and SHAP contribution, while `abs_rt_delta` shows a negative relationship, with larger retention time deviations penalizing the rescoring score. **(b).** XGBoost model feature importance ranked by mean signed SHAP value across top ten XPECTRA input features, confirming `num_matched_ions_total` and `num_matched_y_ions` as the dominant predictors, followed by `neg_log_specEvalue`.

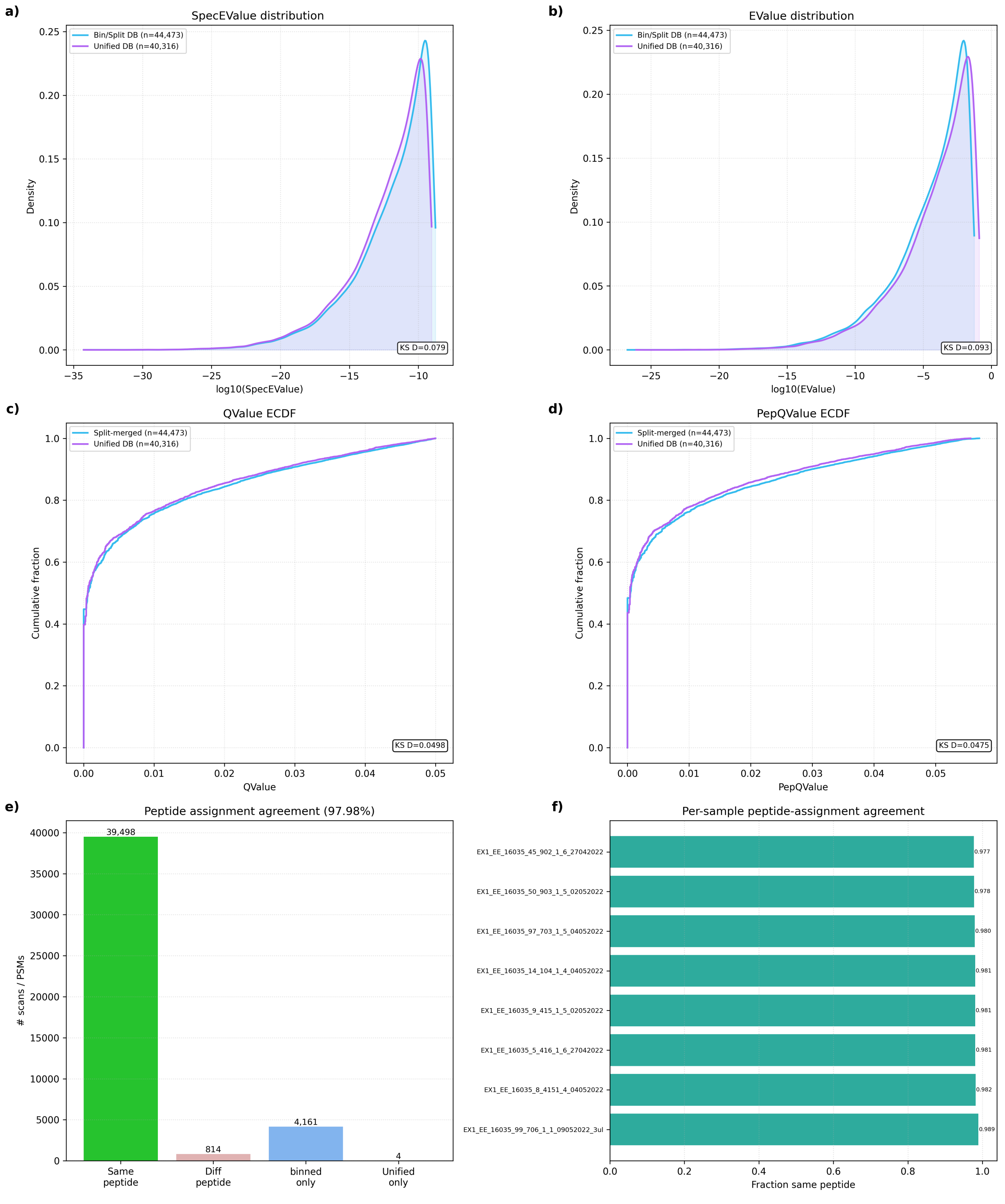


**Fig. S5.** Binning does not bias identification outcomes relative to a unified database search. **(a, b)** Kernel density estimates of $\log_{10}$(SpecEValue) (**a**). and $\log_{10}$(EValue) (**b**). distributions for PSMs identified using the binned/split database (cyan) versus the unified database (purple), showing near-identical score distributions between the two approaches. **(c, d)** Empirical cumulative distribution functions of QValue (**c**). and PepQValue (**d**). for the binned/split (cyan) versus unified (purple) database searches. The maximum empirical differences were small (QValue: $D = 0.0498$; PepQValue: $D = 0.0475$), indicating that the practical distributional shift was limited given the large PSM counts involved. **(e)**. Peptide-level assignment agreement between the binned and unified approaches: the large majority of scans/PSMs (48,467) are assigned the same peptide in both approaches, with a minority assigned different peptides (1,086) or identified uniquely by one approach (2,723 split-only; 2,729 unified-only), yielding an overall peptide-assignment agreement of 97.81%. **(f)**. Per-sample peptide-assignment agreement across eight individual samples from PXD057701, showing consistently high agreement (fraction same peptide range: 0.974–0.985) across samples. All results shown reflect PSMs after XPECTRA rescoring.

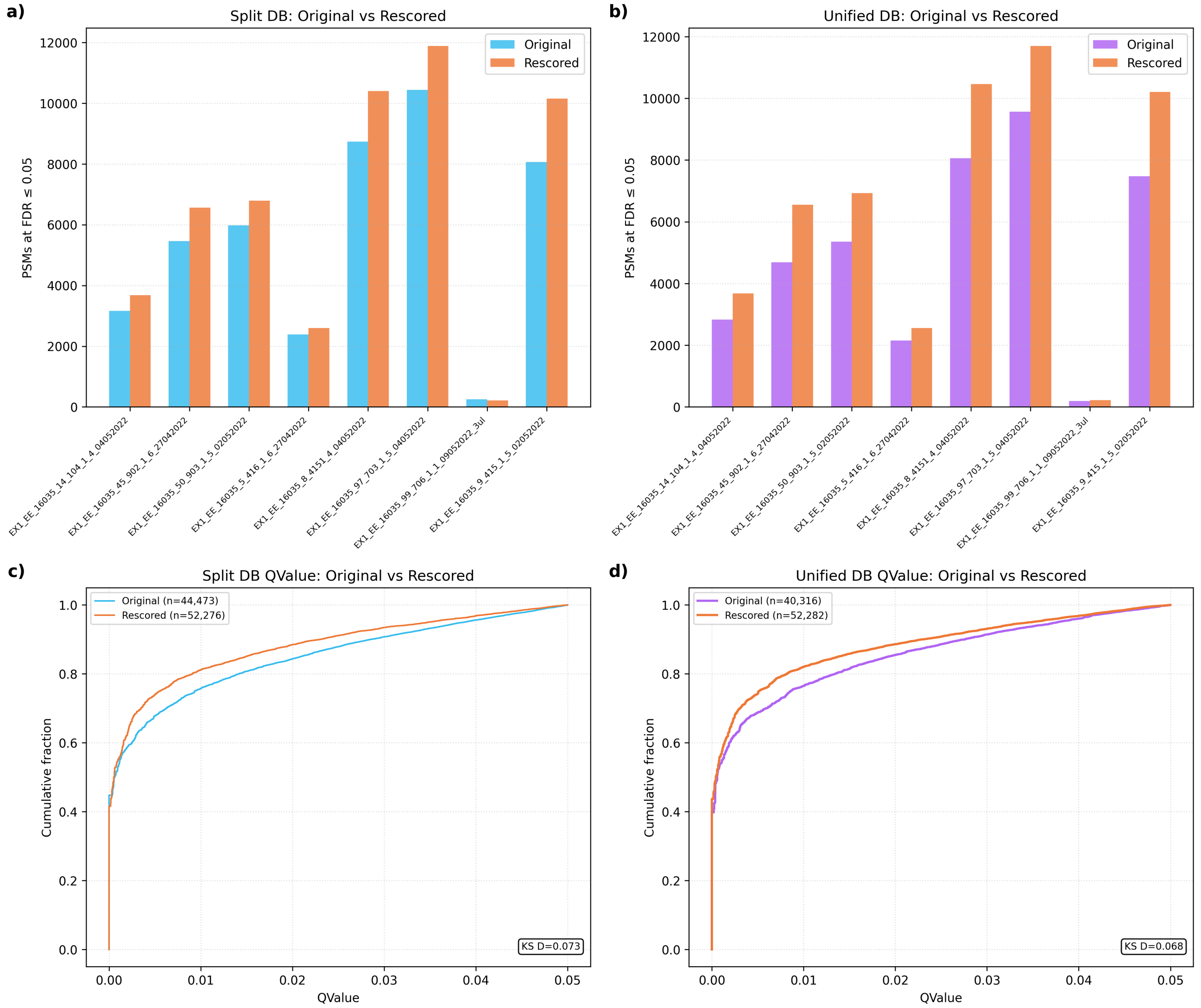


**Fig. S6.** XPECTRA rescoring increases PSM identifications with improved FDR on both binned and Unified database searches. **(a)** Number of PSMs identified at FDR $\leq$ 0.05 before (Original) and after (Rescored) XPECTRA rescoring, per run, for the binned (split) database search strategy. **(b)** Same comparison for the unified database search strategy. **(c)** Empirical cumulative distribution of PSM Q-values (0–0.05 range) before and after rescoring for the split database strategy, pooled across all runs (Original: $n$ = 44,473; Rescored: $n$ = 52,276). The Kolmogorov–Smirnov statistic (KS $D = 0.073$) quantifies the shift toward lower Q-values after rescoring. **(d)** Same comparison for the unified database strategy (Original: $n = 40,316$; Rescored: $n = 52,282$; KS $D = 0.068$). Across both database strategies, XPECTRA rescoring increased PSM yield at FDR $\leq 0.05$ in every run and shifted the Q-value distribution toward higher confidence, indicating consistent rescoring gains independent of database construction strategy.

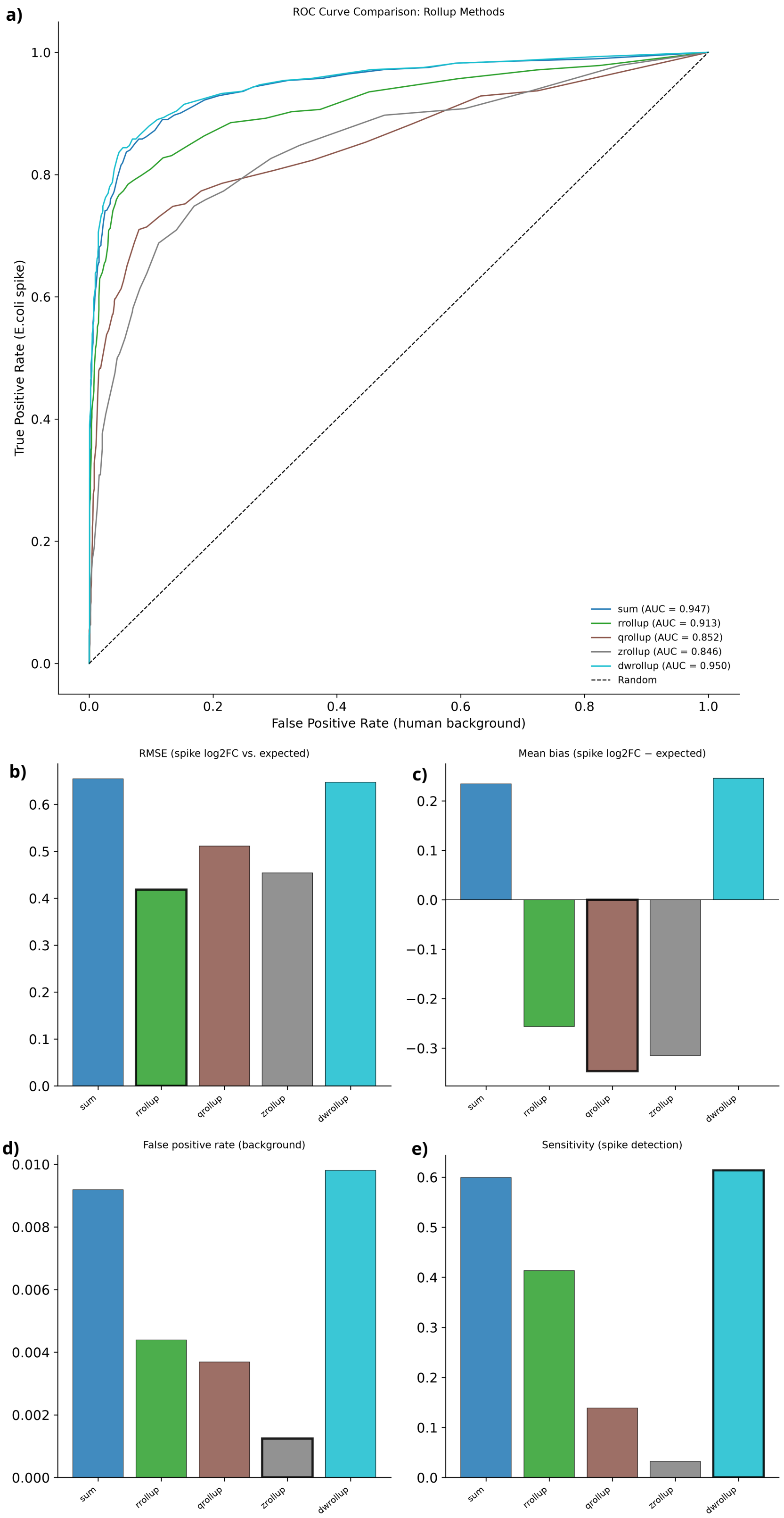


**Fig. S7.** Rollup method choice involves trade-offs across error, bias, and sensitivity, with `DWrollup` achieving the best overall discrimination (AUC). **(a)** ROC curves discriminating *E. coli* spike-in peptides (true positives) from the human background (false positives) for five protein rollup strategies: sum, rrollup, qrollup, zrollup, and weighted (AUC = 0.942, 0.928, 0.857, 0.849, and 0.944, respectively), compared to random classification (dashed diagonal). **(b)** Root-mean-square error (RMSE) between observed and expected $\log_2$ fold-change for spike-in proteins, by rollup method. **(c)** Mean bias (observed − expected $\log_2$ fold-change) by rollup method; values near zero indicate unbiased fold-change estimation. **(d)** False positive rate for background (non-spike) proteins called as differentially abundant, by rollup method. **(e)** Sensitivity for detecting true spike-in proteins as differentially abundant, by rollup method. In panels (b)–(e), the best-performing method for each metric is outlined in black. The weighted rollup achieves the highest AUC, lowest false positive rate among top performers, and highest sensitivity, while maintaining RMSE and bias comparable to the sum method.

### XPECTRA – The rescoring model accompanying MSGFLEX

Tulasi Rao Relangi[1], Harrison A. Hall[1], Aaron T. Wright[1*]

[1*]Department of Biology, Baylor University, Waco, TX, USA.

#### Supplementary Methods

XPECTRA rescoring integrates sequence, spectral, and retention-time evidence within a cross-validated XGBoost ensemble and combines the resulting predictions with native MS-GF+ spectral evidence. All feature transformations and empirically derived quantities, including peptide-level score statistics, retention-time calibration parameters, and spectral-score percentile references, are estimated exclusively from the training data of each cross-validation fold and transferred unchanged to the corresponding held-out partition to prevent information leakage.

#### XPECTRA Feature Construction and Calibration

The four evidence categories introduced in the main-text XPECTRA methods section are constructed as follows.

*Selected MS-GF+ search features.* Log-transformed MS-GF+ score, $-\log_{10}$(spectral E-value), log *de novo* score, and the spectral E-value's percentile rank against a charge-$\times$ length-binned reference distribution computed from the training fold only.

*Tandem-mass-spectral evidence.* Precursor and fragment mass error (signed and absolute, ppm and Da), chromatographic peak width and area, *b-/y-ion* coverage (minimum, maximum, ratio, and balance between the two series), longest *b-/y-ion* series length ratio, mean fragment mass error, precursor-to-fragment error ratio, and within-scan competitive features — the score gap to the runner-up PSM at the same spectrum (raw and normalized), and the number of competing PSMs at that spectrum — which quantify how unambiguous the top match is relative to its nearest competitor.

*Retention-time evidence.* The absolute deviation between predicted and observed retention time, and a retention-time match likelihood; both are outputs of the calibration procedure detailed in Retention-time prediction and calibration section.

*Peptide- and precursor-level characteristics.* Peptide length, charge-to-length ratio, basic/acidic residue counts, hydrophobic residue fraction, N-/C-terminal residue hydrophobicity, C-terminal-half hydrophobic fraction, missed-cleavage count, tryptic-boundary flags (N-terminal basic residue, C-terminal K/R, presence of a KP/RP motif blocking tryptic cleavage), charge × length and charge × mass-error interaction terms, and training-fold-derived peptide observation statistics (observation count, best and median spectral score, score standard deviation, and reliability metric defined as best score normalized by observation count,

$$\text{reliability}(p) = \frac{\text{score}_{\text{best}}(p)}{n_{\text{obs}}(p)}, \tag{S1}$$

which favors peptides identified repeatedly at high confidence over singleton high-scoring hits). The within-scan delta-score feature is normalized against the top-ranked spectral score in the same spectrum,

$$\Delta_{\text{norm}} = \text{clip}\left(\frac{\text{score}_{r1} - \text{score}_{r2}}{|\text{score}_{r1}| + \epsilon},\ 0,\ 1\right), \tag{S2}$$

where $\text{score}_{r1}$ and $\text{score}_{r2}$ are the top- and second-ranked spectral scores at a given spectrum and $\epsilon$ is a small constant preventing division by zero for near-zero top scores.

The native spectral evidence term $e$ used in the main-text fusion equation (Eq. 2 of main Methods Section) is obtained by an auto-calibrated exponential transform of $-\log_{10}(\text{spectral E-value})$,

$$e = 1 - \exp(-s \cdot x), \qquad s = \frac{\ln 2}{\text{median}(x)}, \tag{S3}$$

where $x = -\log_{10}(\text{spectral E-value})$ and the scale $s$ is set, per training fold, so that the median value of $x$ maps to $e = 0.5$; this keeps $e$ on a comparable

$$0, 1$$

scale to the ensemble probability $p$ regardless of the native E-value range of a given dataset.

*Isotope-pattern features.* Two precursor-isotope features were derived from the MS1 survey spectrum. An isotope-pattern consistency score quantified agreement between the observed and averagine-expected isotope envelope[1],

$$S_{\text{iso}} = \max\left(0,\, 1 - \left| \frac{I_{M+1}}{I_M + \varepsilon} - \min(0.0011\, m_{\text{pep}},\, 0.8) \right| \right), \tag{S4}$$

where $I_M$ and $I_{M+1}$ denote the monoisotopic and first isotopic peak intensities, respectively. A binary chimeric-precursor indicator was defined as

$$C = \begin{cases} 1, & \dfrac{I_{M+2}}{I_{M+1} + \varepsilon} > 1.5, \\ 0, & \text{otherwise.} \end{cases} \tag{S5}$$

Higher values of $S_{\text{iso}}$ indicate closer agreement with the expected isotope envelope, whereas $C = 1$ indicates a potentially contaminated precursor isolation window.

All feature transformations and training-fold statistics referenced above are, as stated in the main text, fit only within the training partition of each cross-validation fold and applied unchanged to the held-out fold.

#### Retention-time prediction and calibration

Retention time is predicted in two stages from peptide sequence alone, then calibrated against training-fold observed retention times.

*Base hydrophobicity score.* An SSRCalc-style retention coefficient score is computed using the Krokhin 300 Å reversed-phase retention coefficient set[2], with a peptide-length correction and an N-terminal bonus term:

$$H(p) = \sum_{a \in p} RC_{300\text{Å}}(a) \; - \; 0.35 \ln(1 + L_p) \; + \; 0.42\, RC_{300\text{Å}}(a_1), \tag{S6}$$

where $L_p$ is peptide length and $a_1$ the N-terminal residue. Because raw hydrophobicity scores are not linearly related to observed retention time across a chromatographic gradient, $H(p)$ serves only as a monotonic anchor for the calibration that follows.

*Stage 1 — isotonic warp and local regression.* Within each training fold, target PSMs with valid observed retention time are first corrected for modification-induced retention shifts, then used to fit a monotonic isotonic regression from $H(p)$ onto a LOWESS-smoothed observed-retention-time target, correcting gradient-specific non-linearity while preserving elution order. The warped scores are refit with a second LOWESS pass after interquartile-range outlier removal, and a bias-correction term — the median residual between the LOWESS fit and the full, uncleaned training set — is added so predictions remain centered on the true data distribution rather than the outlier-filtered subset used for smoothing:

$$\text{RT}_{\text{stage1}}(p) = \text{LOWESS}\big(\text{warp}(H(p))\big) + \delta_{\text{bias}}. \tag{S7}$$

*Stage 2 — residual regression.* The systematic residual left by Stage 1, $r(p) = \mathrm{RT}_{\mathrm{obs}} - \mathrm{RT}_{\mathrm{stage1}}(p)$, is modeled with a gradient-boosted regressor (XGBoost, absolute-error objective) trained on the warped hydrophobicity anchor, peptide length and its square, charge and its interaction with length, fractional amino-acid composition, N-/C-terminal residue identity, Kyte–Doolittle hydrophobicity summary statistics (global and positional), and modification summary features. Sample weights combine a rank-based spectral-confidence term with soft down-weighting of extreme residuals so that outlier retention-time observations do not dominate the fit; regressor capacity scales with training-fold size.

*Final prediction and match score.* The calibrated retention-time prediction is:

$$\mathrm{RT}_{\mathrm{pred}}(p) = \mathrm{RT}_{\mathrm{stage1}}(p) + \hat{r}_{\mathrm{XGB}}(p), \tag{S8}$$

and the retention-time match feature supplied to XPECTRA is a Laplace likelihood,

$$\mathrm{rt\_score}(p) = \exp\left(-\frac{|\mathrm{RT}_{\mathrm{obs}} - \mathrm{RT}_{\mathrm{pred}}(p)|}{\mathrm{MAD}_{c,\ell}}\right), \tag{S9}$$

where $\mathrm{MAD}_{c,\ell}$ is the median absolute residual deviation stratified by charge state $c$ and peptide-length bin $\ell$, falling back to a single training-fold MAD where a given stratum has insufficient support. This stratification reflects that retention-time prediction error is not homoscedastic across charge state and peptide length. Across all seven benchmarked datasets, the calibrator achieves a normalized mean absolute error (MAE divided by the observed retention-time range) below 0.1.

#### Automatic modification detection and correction

Modification handling requires no per-dataset manual annotation. Post-translational modification and TMT-labeling status are parsed directly from each peptide's PHRP-notation modification string at run time, so every characterized modification instance — oxidation, phosphorylation, deamidation, acetylation, and the TMT-loss variant AcNoTMT — is automatically detected, counted, and localized per peptide without any dataset-specific configuration step. These automatically extracted counts populate the modification-derived features in the main feature vector (total modification count, cumulative modification mass, mass per residue, and per-type counts) and feed the retention-time correction below.

Retention-time offsets attributable to each modification type are likewise learned automatically rather than fixed. For each training fold, matched pairs of modified and unmodified forms of the same core peptide sequence are identified, and a Huber-regularized linear model is fit relating modification count to the observed retention-time shift:

$$\Delta \mathrm{RT}_m = \omega_m \cdot n_m(p) + \eta, \qquad \{\omega_m, \eta\} = \arg\min_{\omega,\eta} \sum_i \ell_{\mathrm{Huber}}\big(\Delta \mathrm{RT}_{m,i} - \omega\, n_m(p_i) - \eta\big), \tag{S10}$$

where $n_m(p)$ is the count of modification type $m$ on peptide $p$ and $\ell_{\mathrm{Huber}}$ is the Huber loss, giving robustness to outlier pairs. Each fitted offset $\omega_m$ is checked against a physically plausible range (for example, phosphorylation is constrained to shift retention earlier, within an empirically bounded window); an offset outside its plausible range is automatically rejected in favor of a literature-informed default. This per-fold, per-dataset re-estimation allows the same calibration procedure to adapt to differing modification prevalence and chromatographic conditions across datasets without any code or configuration changes.

##### Cross-validation and ensemble configuration

As described in the main text, PSMs are partitioned into five group-stratified folds by `OptimalScanNumber` (stratified by target/decoy status) to prevent within-spectrum leakage between competing candidate matches for the same spectrum. Within each fold, five independently seeded XGBoost classifiers are trained on the remaining folds using hyperparameters selected automatically by training-fold size, with a stratified 10% internal holdout used for early stopping. Raw classifier margins are converted to probabilities via the sigmoid function, $\sigma(z) = 1/(1 + e^{-z})$, and the five seed-level predictions per PSM are combined by trimmed mean to yield the fold's out-of-fold `rescore_prob`, reducing sensitivity to any single model initialization.

$$\texttt{rescore\_prob}(p) = \frac{1}{3} \sum_{k=2}^{4} \sigma\big(z_{(k)}(p)\big), \tag{S11}$$

where $z_{(1)} \leq \cdots \leq z_{(5)}$ are the five seed-level classifier margins sorted in ascending order and the lowest and highest are discarded before averaging, reducing sensitivity to any single model initialization.
